# Accounting for Edge Uncertainty in Stochastic Actor-Oriented Models for Dynamic Network Analysis

**DOI:** 10.1101/2025.02.17.638664

**Authors:** Heather M. Shappell, Mark A. Kramer, Catherine J. Chu, Eric D. Kolaczyk

## Abstract

Stochastic Actor-Oriented Models (SAOMs) were designed in the social network setting to capture network dynamics representing a variety of influences on network change. The standard framework assumes the observed networks are free of false positive and false negative edges, which may be an unrealistic assumption. We propose a hidden Markov model (HMM) extension to these models, consisting of two components: 1) a latent model, which assumes that the unobserved, true networks evolve according to a Markov process as they do in the SAOM framework; and 2) a measurement model, which describes the conditional distribution of the observed networks given the true networks. An expectation-maximization algorithm is developed for parameter estimation. We address the computational challenge posed by a massive discrete state space, of a size exponentially increasing in the number of vertices, through the use of the missing information principle and particle filtering. We present results from a simulation study, demonstrating our approach offers improvement in accuracy of estimation, in contrast to the standard SAOM, when the underlying networks are observed with noise. We apply our method to functional brain networks inferred from electroencephalogram data, revealing larger effect sizes when compared to the naive approach of fitting the standard SAOM.

## 1 Introduction

Networks have been used broadly in biology, the social sciences, and many other fields to model and analyze the relational structure of individual units in a complex system. In a network model, nodes or vertices represent the units of the system, and edges connect the vertices if the corresponding units share a relationship. It is desirable, in many applications, to study the change in connections among the vertices of a network over time. Stochastic actor-oriented models (SAOMs), designed specifically for longitudinally observed networks (i.e. network panel data), are a class of models that were developed for this purpose in the social network setting by Snijders et al. [33, 37, 34].

The SAOM framework revolves around the notion that the vertices control the connections they make to other vertices. This approach is different from other models, such as the temporal exponential random graph model (TERGM) developed by Hanneke et al. [17]. The assumption is that the network evolves as a continuous time Markov chain and that the networks one has observed are snapshots of this stochastic process. Network changes are assumed to happen by one vertex making a change in one of its connections at a time. Vertices seek to change these connections such that their ‘personal satisfaction’ with the network configuration is maximized. This ‘satisfaction’ is captured by an objective function, in the form of a linear combination of effects, which can be both endogenous (i.e. functions of the network itself) and exogenous (i.e. functions of vertex characteristics). Parameters indicating the strength of each effect are estimated using either a method of moments or maximum likelihood simulation-based approach, and hypotheses associated with each effect can be tested, similar to a linear regression framework [34, 35].

SAOMs have been shown to be useful in a variety of applications, but a key assumption is that the networks one has observed are error-free. In other words, one is assuming that the vertices present in the network and the relationships observed among them are all accurate at the time the network data were measured. However, what if this is not the case? Network analysis has long been plagued by issues of measurement error [43]. For instance, survey respondents may not report the correct spellings of their friends’ names. This not only leads to erroneous vertices, but also to an absence of an edge to the correct vertex in the social network. Furthermore, even if everyone reports the correct spellings of their friends’ names, the understanding of what qualifies as a friendship tie can vary by respondent. Other settings, such as co-authorship networks, which represent collaborative relationships, can also contain false positive and false negative edges [44]. In this case, false edges may exist because of failure to account for edge decay. One can deal with this issue by setting a pre-specified time window under which the established relationship is thought to be meaningful [11]. However, setting too narrow of a window might overlook important relationships and introduce false negatives, while setting too wide of a window can introduce false positives [43].

In addition to SAOMs being used extensively in a social network context, in our own work we have recently adapted these models to resting-state fMRI complex brain networks. We sought to answer questions such as, “If two brain regions are in the same cortical lobe, are they more likely to connect?” In this case, a connection represents a similar pattern of brain activity. In the neuroscience setting, functional network edges are almost always defined based on some measure of association between patterns of activation between distinct brain regions [31]. Therefore, it is unreasonable to assume that the observed networks are always the truth. Instead, we expect some level of type I error (false positives) and type 2 error (false negatives) to exist in the inferred networks.

Motivated by scenarios such as these, our goal is to account for false positive and false negative edges while analyzing observed networks with SAOMs. To capture the notion of false positive and false negative rates, along with the parameters in the SAOM, we propose a hidden Markov model (HMM) based approach. This modeling approach consists of two components - the latent Markov model and the measurement model. The latent Markov model specifies that the unobserved hidden networks evolve according to a Markov process, as they did in the original SAOM framework. The measurement model describes the conditional distribution of the observed networks given the true networks.

HMMs, developed by Baum and colleagues in the 1960s [2], are a natural modeling approach to take given that we have an observable sequence of a system in which the hidden state is governed by a Markov process. They have been widely studied in statistics [8] and have been applied extensively in many applications, such as in speech recognition and biological sequence analysis [12, 46]. While HMMs have been used much less frequently in the dynamic network analysis literature, there is some work where they have been incorporated. For example, Guo et al. developed the hidden TERGM, which utilizes a hidden Markov process to model and recover temporally rewiring networks from time series of node characteristics [16]. Dong et al. present the Graph-Coupled Hidden Markov Model, a discrete-time model for analyzing the interactions between individuals in a dynamic social network [6]. Their method incorporates dynamic social network structure into a hidden Markov Model to predict how the spread of illness occurs and can be avoided on an individual level. Similarly, Raghavan et al. propose a coupled Hidden Markov Model, where each user’s activity in a social network evolves according to a Markov chain with a hidden state that is influenced by the collective activity of the friends of the user [24].

HMMs for SAOMs have not been published in a peer-reviewed journal, but they have been published in a dissertation [20]. Lospinoso worked to tackle this same problem of accounting for error on the observed networking edges in the SAOM setting. We take a similar approach in this paper, but instead of using a full MCMC algorithm for performing maximum likelihood estimation of the model parameters, we take an MCMC within an Expectation-Maximization approach to parameter estimation. We also focus on a brain network application, whereas Lospinoso applies the methodology to social networks. We touch further on the similarities and differences to Lospinoso’s approach in the Discussion section.

The remainder of this paper is organized as follows. In section 2, we introduce our HMM-SAOM set-up. Section 3 provides necessary background information on the SAOM model set-up, as well as the estimation routine traditionally used to fit the SAOM parameters. Our framework incorporates the methods described in this section. Section 4 describes our Expectation-Maximum algorithm for maximum likelihood estimation of the false positive and false negative error rates, along with the SAOM parameters. We assess the performance of our method on a series of simulated dynamic networks in Section 5, comparing it to the case of fitting only a standard SAOM to noisy networks. In Section 6, we apply our method to functional brain networks inferred from electroencephalogram (EEG) data. We conclude with a discussion of our method and open directions for future research.

## 2 The SAOM Hidden Markov Model Set-Up

We consider repeated observations of a directed network on a given set of vertices 𝒩 = 1, …, *N*, observed according to a panel design. The observations are represented as a sequence of digraphs *y*(*t*_*m*_) for *m*= 1, …, *M*, where *t*_1_ < … < *t*_*M*_ are the observation times and the node set is the same for all observation times. A digraph is defined as a subset *y* of {(*i, j*) ∈ 𝒩^2^|*i* ≠ *j*} [35, 34]. When (*i, j*) ∈ *y*, there is an edge from vertex *i* to vertex *j*. The observations *y*(*t*_*m*_) are realizations of random digraph/network variables *Y (t*_*m*_). This random vector of observed network variables *Y* (*t*_1_), …, *Y* (*t*_*M*_) is denoted by 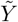. We represent a true/hidden network variable underlying the observed network at a particular observation time by *U*(*t*_*m*_). The vector of true network variables, *U*(*t*_1_), …, *U*(*t*_*M*_) is denoted by *Ũ*. See Figure 1 for a visual representation. We also assume that

**Figure 1:**
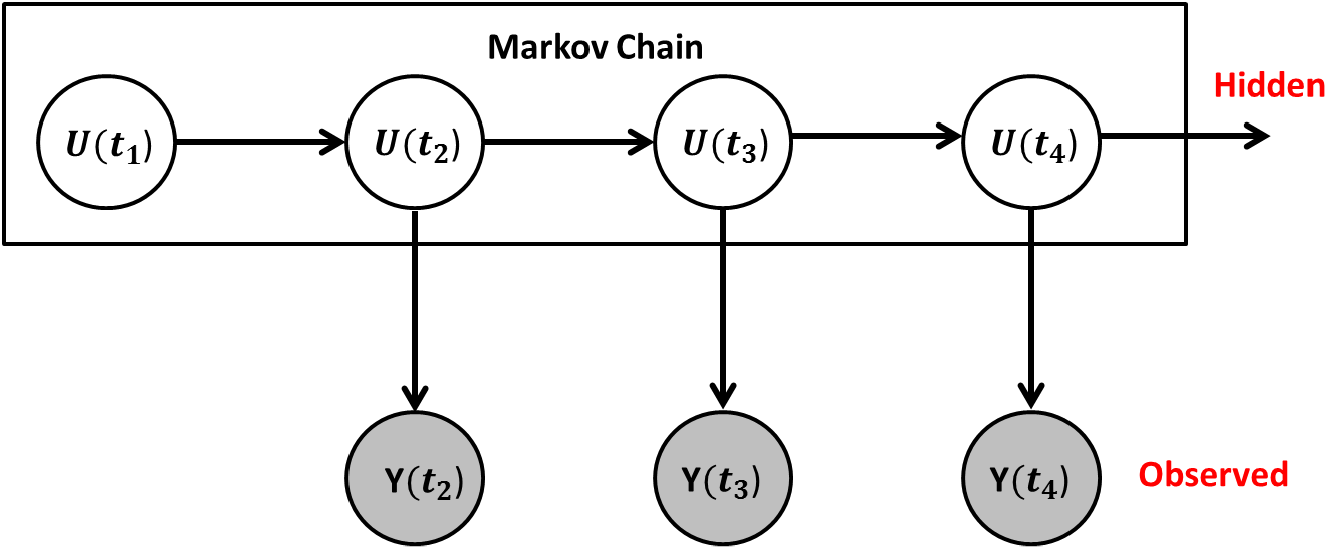
Hidden Markov Model Set-Up. The unobserved hidden networks evolve according to a Markov process, as they did in the original SAOM framework. The true networks are then observed with measurement error.

1. The vector of true networks follows a first order Markov chain.

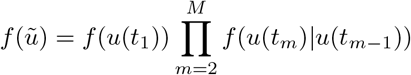
2. The observed networks are conditionally independent given the latent process.

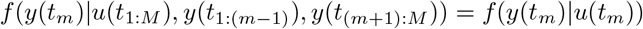
3. We condition on the first observed network *y*(*t*_1_), and we assume the first true network *u*(*t*_1_) is observed error-free.

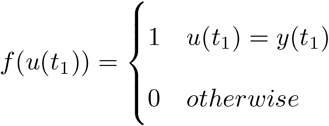

Therefore, the complete data log-likelihood, conditional on *y*(*t*_1_), can be written as:

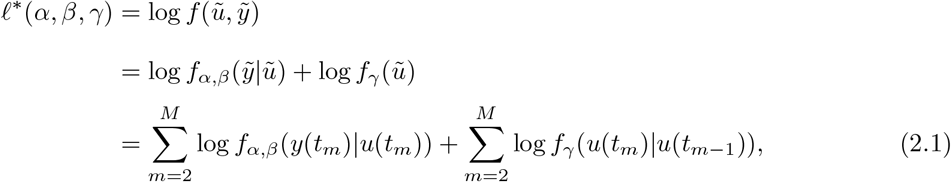

where *α* is the false positive rate, *β* is the false negative rate, and *γ* consists of the objective function parameters and rate parameters in the SAOM. Additional details on the SAOM and its parameters are provided in Section 3.

The first term in *l*\* (*α, β, γ*) derives from the conditional distribution of the observed networks given the true networks and takes the form

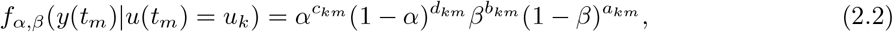

where *a*_*km*_, *b*_*km*_, *c*_*km*_, and *d*_*km*_ (for a given true network *u*_*k*_ at observation time *t*_*m*_) represent counts of false positive and false negative edges (see Table 1).

**Table 1:** Counts Corresponding to False Positive and False Negative Edges.

| True/Observed | Edges | Non-Edges |
| --- | --- | --- |
| Edges | a | b |
| Non-Edges | c | d |

The second term in *l*\* (*α, β, γ*), deriving from the network transition probability distribution, cannot be calculated in closed form. Additional details will be provided in the Section 3.

## 3 Background on SAOM framework

### 3.1 Modeling framework

We adopt the SAOM framework of Snijders et al. [37] for the evolution of our true networks, which are underlying the sequence of our observed networks, *y*(*t*_*m*_). In this framework, it is assumed that the changing network is the outcome of a continuous time Markov process with time parameter *t* ∈ *T* where the *u*(*t*_*m*_), underlying the *y*(*t*_*m*_), are realizations of stochastic digraphs *U*(*t*_*m*_) embedded in a continuous-time stochastic process *U*(*t*), *t*_1_ ≤ *t* ≤ *t*_*M*_. The totality of possible networks is the state space and the discrete set of true networks are snapshots of the true network state during the continuous period of time. In other words, many changes are assumed to happen in the true networks between observation times, and the process unfolds in time steps of potentially varying lengths [33, 37, 34].

Each *U*(*t*) is made up of *N* × (*N* − 1) possible edge status variables *u*_*ij*_, where *u*_*ij*_ = 1 if there exists a directed edge from vertex *i* to vertex *j* and *u*_*ij*_ = 0 otherwise. At a given moment, one probabilistically selected vertex may change an edge, where the decision is modeled according to a random utility model, requiring the specification of a utility function (i.e., objective function) depending on a set of explanatory variables and parameters. Therefore, we are reduced to modeling the change of one edge status variable *u*_*ij*_ by one vertex at a time (a network micro step) and modeling the occurrence of all these micro steps over time. The first true network *u*(*t*_1_) serves as a starting value of the evolution process. At any time point *t* with current network *u*(*t*) = *u*, each of the vertices has a rate function *λ*_*i*_(*δ, u*), where *δ* is a parameter. Therefore, the waiting time until occurrence of the next micro step by any vertex is exponentially distributed with parameter

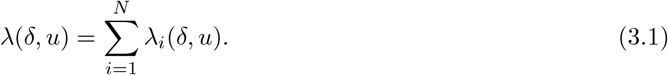

Given that an opportunity for change occurs, the probability that it is vertex *i* who gets the opportunity is given by

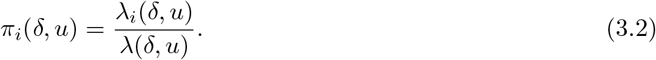

The micro step that vertex *i* takes is determined probabilistically by a linear combination of effects. For example, let’s assume that *u* is the current network and vertex *i* has the opportunity to make a network change. The next network state *u*^′^ then must be either equal to *u* or deviate from *u* by one edge. Vertex *i* chooses the value of *u*^′^ for which

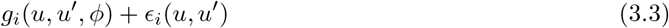

is maximal, where *ϵ*_*i*_(*u, u*^′^) is a Gumbel-distributed random disturbance that captures the uncertainty stemming from unknown factors, and

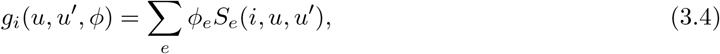

where *ϕ*_*e*_ represent parameters and *S*_*e*_(*i, u, u*^′^) represent the corresponding effects. There are many types of effects one can place in the model. See Ripley et al. [26] for a full list. Some are purely structural effects, such as triangle formation and reciprocity. Other effects may involve vertex traits, such as gender or smoking status of the individuals in a social network.

Equation 3.4 is the objective function. It can be thought of as a function of the network perceived by the focal vertex. Probabilities are higher for moving towards network states with a high value of the objective function. The objective function depends on the personal network position of vertex *i*, vertex *i*’s exogenous covariates, and the exogenous covariates of all of the vertices in *i*’s personal network. Due to distributional assumptions placed on *ϵ*_*i*_(*u, u*^′^), the probability of choosing *u*^′^ can be expressed in multinomial logit form as

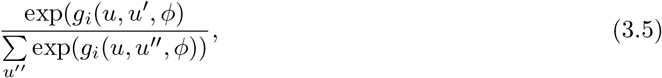

where the sum of the denominator extends over all possible next network states *u*^′′^.

For each set of model parameters, there exists a stationary distribution of probabilities over the state space of all possible network configurations for the Markov process that governs the SAOM. The complexity of the model does not allow for the equilibrium distribution (nor the likelihood of the network ‘snapshots’) to be calculated in closed form. Therefore, parameter estimates need to be obtained via an iterative stochastic approximation version of a maximum likelihood approach based on data augmentation.

### 3.2 SAOM maximum likelihood framework

The distribution of the true networks *U*(*t*_1_), …, *U*(*t*_*M*_) conditional on *U*(*t*_1_) cannot generally be expressed in closed form. Therefore, the true networks are augmented with data such that an easily computable likelihood is obtained [35]. The data augmentation can be done for each period (*U*(*t*_*m* − 1_), *U*(*t*_*m*_)) separately, and therefore, it is explained below only for *U*(*t*_1_) and *U*(*t*_2_).

Denote the time points of an opportunity for change by *T*_*r*_ and their total number between *t*_1_ and *t*_2_ by *R*, the time points being ordered increasingly so that *t*_1_ *= T*_0_ *< T*_1_ *< T*_2_ *<* … *< T*_*R*_ ≤ *t*_2_. The model assumptions imply that at each time *T*_*r*_, there is one vertex, denoted *I*_*r*_, who gets an opportunity for change at this time moment. Define *J*_*r*_ as the vertex toward whom the edge status variable is changed, and define *J*_*r*_ = *I*_*r*_ if there is no change. Given *u*(*t*_1_), the outcome of the stochastic process (*T*_*r*_, *I*_*r*_, *J*_*r*_), *r* = 1, …, *R* completely determines *u*(*t*), *t*_1_ < *t* ≤ *t*_2_.

The stochastic process *V* = ((*I*_*r*_, *J*_*r*_), *r* = 1, …, *R*) will be referred to as the sample path. Define *u*^(*r*)^ = *u*(*T*_*r*_). The graphs *u*^(*r*)^ and *u*^(*r* − 1)^ differ in element (*I*_*r*_, *J*_*r*_), provided *I*_*r*_ ≠ *J*_*r*_ and in no other elements. Snijders et al. show that in the case where the vertex-level rates of change *λ*_*i*_(*δ, u*) are constant (which is an assumption typically recommended in using SAOMs), denoted by *δ*_1_, the probability function of the sample path, conditional on *u*(*t*_1_), is given by

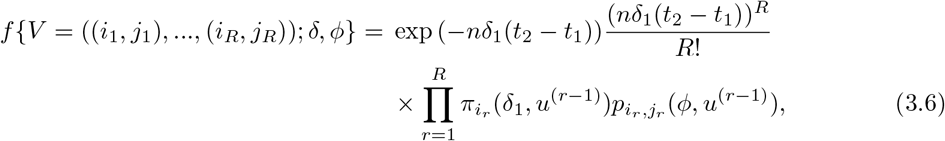

where *π*_*i*_ is defined in (3.2), and

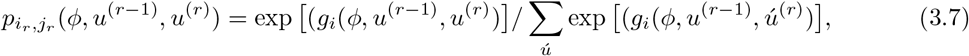

where the summation extends over all possible next network states *ú*.

Therefore, for two possible true networks (*u*(*t*_1_), *u*(*t*_2_)) augmented by a sample path, the likelihood conditional on *u*(*t*_1_) can be expressed exactly. An MCMC algorithm is used to find the maximum likelihood estimator based on the augmented data. The algorithm Snijders et al. implements, proposed by Gu and Kong [15], is based on the missing information principle, which can be summarized as follows. Suppose *ũ* is given and having probability density *f*(*ũ*; *γ*). Then, suppose it is augmented by extra data *v*, such that the *j*oint density is *f*(*ũ, v*; *γ*). Denote the incomplete data score function 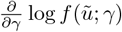 by *S*(*γ*; *ũ*) and the complete data score function 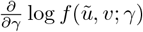 by *S*(*γ*; *ũ, v*). It can be shown

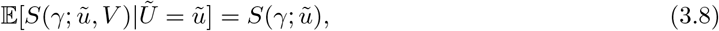

which implies that maximum likelihood estimates can be determined as the solution to

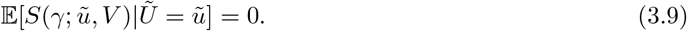

In the SAOM context, *U*(*t*_1_) is treated as fixed, and data are augmented between the true networks at each observed time point *m* = 1, …, *M* by a sample path that could have brought each true network to the next. Each period *(t*_*m* − 1_, *t*_*m*_) is treated separately, and draws from the probability distribution of the sample path, *v*_*m*_, conditional on *U*(*t*_*m*_) = *U*(*t*_*m*_), *u*(*t*_*m* − 1_) = *u*(*t*_*m* − 1_), are generated by the Metropolis Hastings Algorithm. These sample paths for each period combined constitute *v*. Let the rate and objective function parameters (*δ, ϕ*) be denoted by *γ*. The complete data score function can be written as

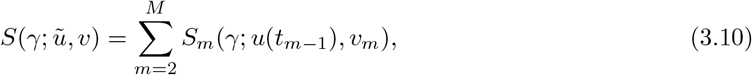

and (3.8) can now be written as

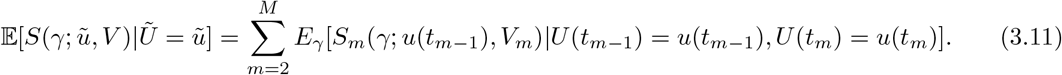

The maximum likelihood estimate is the value of *γ* for which (3.11) equals 0. The solution is obtained by stochastic approximation via a Robbins Monro Algorithm with updating step

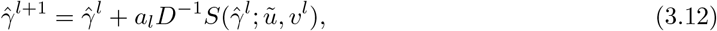

where *v*^*l*^ is generated according to the conditional distribution of *V*, given *Ũ* = *ũ*, with parameter value 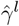. *a*_*l*_ is a sequence of positive numbers tending to 0, and *D* is a positive definite matrix. See Snijders et al.[35] and https://www.stats.ox.ac.uk/~snijders/siena/Siena_algorithms.pdf[stats.ox. ac.uk] for additional details on this algorithm.

## 4 Maximum Likelihood Estimation for the SAOM-HMM

In this section we present an algorithm to calculate the maximum likelihood estimates of the parameters *α, β*, and *γ* in our SAOM-HMM described in Section 2. Let Γ consist of (*α, β, γ*). We develop a variation of the EM algorithm, an iterative method which alternates between performing an expectation (E) step and a maximization (M) step [5]. We first find the expected value of the complete data log-likelihood (Γ) = log 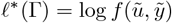, with respect to the unknown, true networks *ũ*, given the observed networks 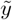 and the current parameter estimates for Γ. We then maximize the expected log-likelihood found in the E-step. Our M-step framework embeds the SAOM estimation routine described in Section 3.2. The overall schematic of our estimation routine *(t*hat we will describe in detail in the remainder of this section) is as follows:

Given a series of observed networks 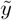 and a number of effects one wishes to include in the SAOM, a SAOM is fit (via the maximum likelihood estimation routine described in Section 3.2) to get initial estimates of the parameters associated with each effect. We also make the choice, for this current implementation, to assume a constant rate parameter across all vertices. This is a fairly standard choice and is recommended in the SAOM literature, unless one has strong reason to believe otherwise. Set 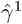 equal to these estimates. The *p*^*th*^ updating step of our EM algorithm then proceeds as follows:

1. Using 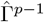, perform Algorithms 1.1, 1.2, and 1.3 described in Section 4.3 to sample a series of true networks. This step corresponds to the E-step in the EM algorithm and is necessary due to the incredibly large state space present in our Expectation.
2. Perform the maximization step using the formulas in section 4.2.1 to obtain 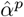 and 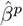.
3. Perform the maximization routine outlined in section 4.2.2 to obtain 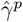.
4. Repeat steps 1-3 until convergence.

Additional details are presented below.

### 4.1 E-Step

The expected value of the complete data log-likelihood (2.1) with respect to the true networks *ũ* given the observed networks 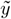 is:

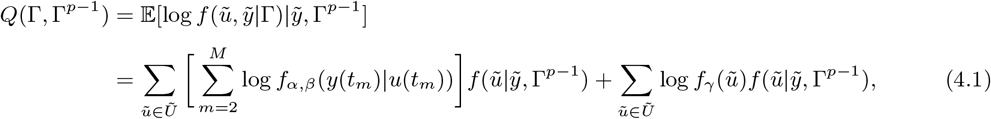

where *p* is the iteration number, Γ^*p* − 1^ are the current parameter estimates that we use to evaluate the expectation, and Γ^*p*^ are the new parameters that we want to optimize to increase *Q*(Γ, Γ^*p* − 1^). This expectation is difficult to calculate due to the magnitude of the state space. In order to address this computational challenge, we employ a particle filtering based sampling scheme (i.e. a sequential Monte Carlo method), which will be described in detail in Section 4.3.

### 4.2 M-Step

The second step of the EM algorithm is to maximize the expectation, i.e. to calculate Γ^*p*^ = arg max_Γ_ *Q*(Γ, Γ^*p* −1^). Since the parameters we wish to optimize are independently split into two terms in *Q*(Γ, Γ^*p* −1^), we can optimize each separately.

#### 4.2.1 Maximizing in *α* and *β*

The first term in *Q*(Γ, Γ^*p* − 1^) takes the form:

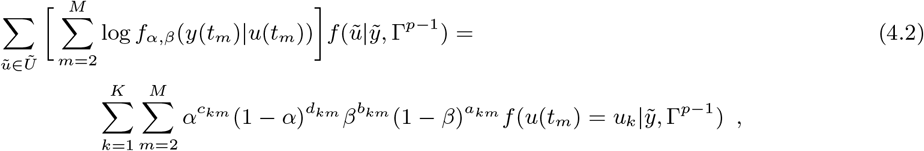

where *a, b, c*, and *d* for a given true network *u*_*k*_ at observation time *t*_*m*_ are defined in Table 1. Let *A, B, C*, and *D* correspond to the random variables for each. Maximizing in *α* and *β* yields the following:

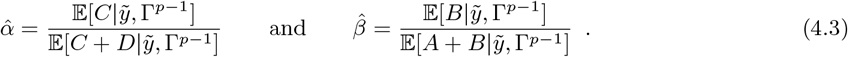

The formula for 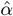 is the expected number of false edges in the observed network divided by the expected number of non-edges in the true network, given the observed networks 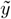 and current parameter estimates Γ^*p* − 1^. The formula for 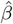 is the expected number of false non-edges in the observed network divided by the expected number of edges in the true network, given the observed networks 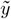 and current parameter estimates Γ^*p* − 1^. The derivation for each of these formulas is provided in Appendix A.

#### 4.2.2 Maximizing in *γ*

The second term in our *Q* function is

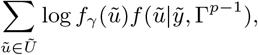

where *γ* consists of the SAOM parameters. We adopt the SAOM framework of Snijders et al. [37] described in Section 2. Note that taking the derivative of the second term in our *Q* function yields

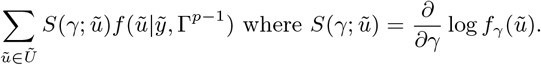

Since *f*_*γ*_ (*ũ*) cannot be calculated in closed form, neither can 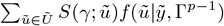. To aid in maximization, we augment each true network series *ũ* with a possible sample path that could have led one true network to the next in the series. We define the random variable *V* to be a sample path associated with a true network series *ũ*. Drawing upon the missing information principle and the work of Snijders et al. [35] and Gu et al. [15], we write the equation above as

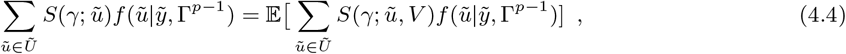

where the expectation on the right-hand side of the equation is with respect to *V*. Maximum likelihood estimates for *γ* can be determined, under regularity conditions, as the solution for (4.4) equaling 15. The proof of (4.4) is shown in Appendix B.

In our approach, the solution to (4.4) is obtained by stochastic approximation via a Robbins Monro algorithm, which is similar to that used in Snijders et al. [35] and described in Section 3.2. At each iteration of the Robbins Monro algorithm, a possible *V*, i.e. a path connecting each possible true network, in each possible latent network series, is sampled. This sampling is done via a Metropolis Hastings algorithm [35]. Next, 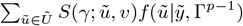is calculated and used in an updating step in the algorithm. It is unreasonable, given the prohibitively large state space of true network series, to sum over every possible true network series in this calculation. Therefore, we note that 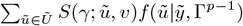 is an expectation, and we sample a smaller number (denoted by *H*) of true network series from 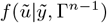. The average of *S*(*γ*; *ũ, v*) for this smaller sample is what is used in the updating step of the Robbins Monro algorithm in place of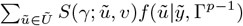. The updating step is

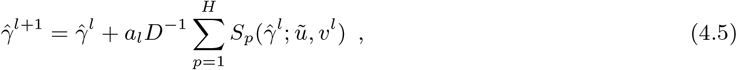

where the sum is over the total number of sampled true network series *ũ, D* is the matrix of partial derivatives, and *a*_*l*_ is a sequence of positive numbers tending to 0.

Our actual implementation of the above algorithm again borrows from that of Snijders et al. [35], which follows directly from the work of Gue and Kong [15]. The algorithm performed consists of two phases. In the first phase a small number of simulations are used to obtain a rough estimate of the matrix of partial derivatives (defined as D in our updating step), which are estimated by a score-function method [29]. The second phase determines the estimate of *γ* by simulating *V* and performing the updating step.

### 4.3 Particle Filtering

The expectation in the E-step of our E-M algorithm is difficult to calculate largely due to the magnitude of the state space. There are 2^*N* * (*N* − 1)^ possible true networks at each observation time, and to calculate 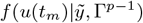 for a given observation moment *m*, one needs to sum over all possible combinations of *u* at the previous *m* − 1 observation times. The forward-backward algorithm can in principle be used to compute these posterior marginals of all hidden state variables given a sequence of observations [23]. The algorithm makes use of the principle of dynamic programming to efficiently compute the values in two passes. The first pass goes forward in time, while the second goes backward in time. However, given the magnitude of our network space, direct use of this approach is not computationally feasible in any but the smallest of problems. Also, as described in the previous section, the transition probabilities, *f*_*γ*_ (*u*(*t*_*m*_)|*u*(*t*_*m* − 1_)), cannot be calculated in closed form.

In order to address this computational challenge, we employ a particle filtering based sampling scheme (i.e. a Sequential Monte Carlo method) [7]. Particle filtering methods provide a way to sample from an approximation of 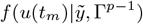through time. Our adaptation of particle filtering principles to the current content is described in Algorithms 1.1 - 1.3 and follows almost exactly from [7]. Algorithm 1.1 mirrors the forward portion of the forward-backward algorithm referred to above. Algorithm 1.2 describes the ancestral simulation of ⋆ in 1.1, and Algorithm 1.3 is used to sample full sequences of latent network variables, i.e., to obtain an approximate sample from the conditional distribution of the latent networks given the observed networks 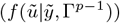. For more details on the particle filtering method we have applied in our algorithm, refer to [7].

#### Algorithm 1.1

A mirror of the Forward Algorithm

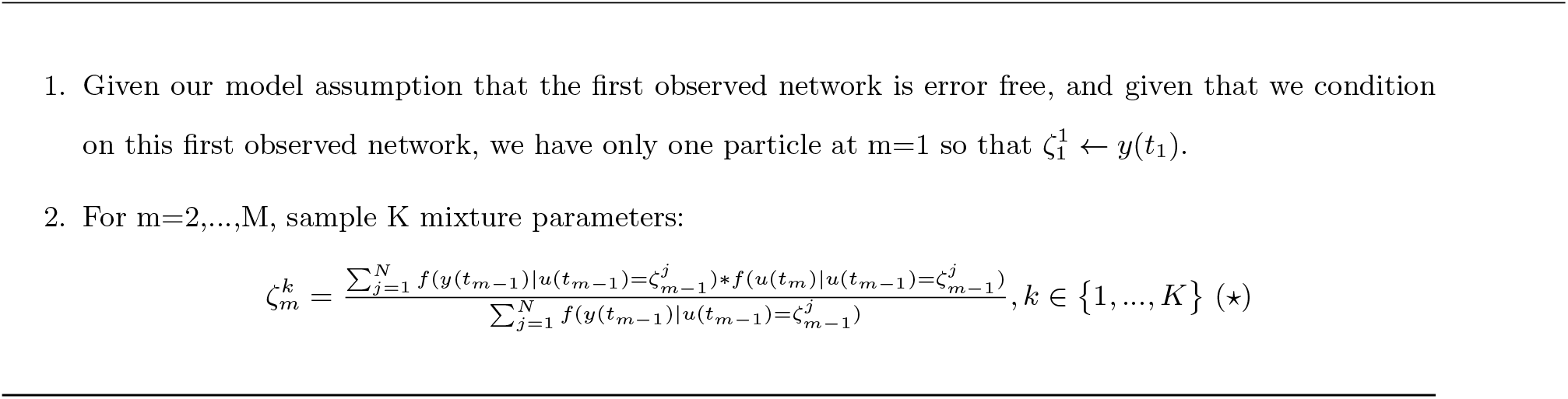

Of special note is that (⋆) in step 2 is sampled in two stages. The first stage samples a mixture parameter, 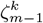, with probability proportional to *f*(*y*(*t*_*m* − 1_)|*u*(*t*_*m* − 1_) = *ζ*_*m* − 1_). The second stage samples from *f*(*u*(*t*_*m*_)|*u*(*t*_*m* − 1_) = *ζ*_*m* − 1)_. This two-stage process provides a genealogical interpretation of the particles that are produced by the algorithm and is outlined in Algorithm 1.2. For *m* ∈ {1, …, *M*} and *k* ∈{1, …, *K*}, we denote by 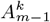 the ancestor index of particle 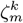.

#### Algorithm 1.2

Ancestral simulation of (⋆)

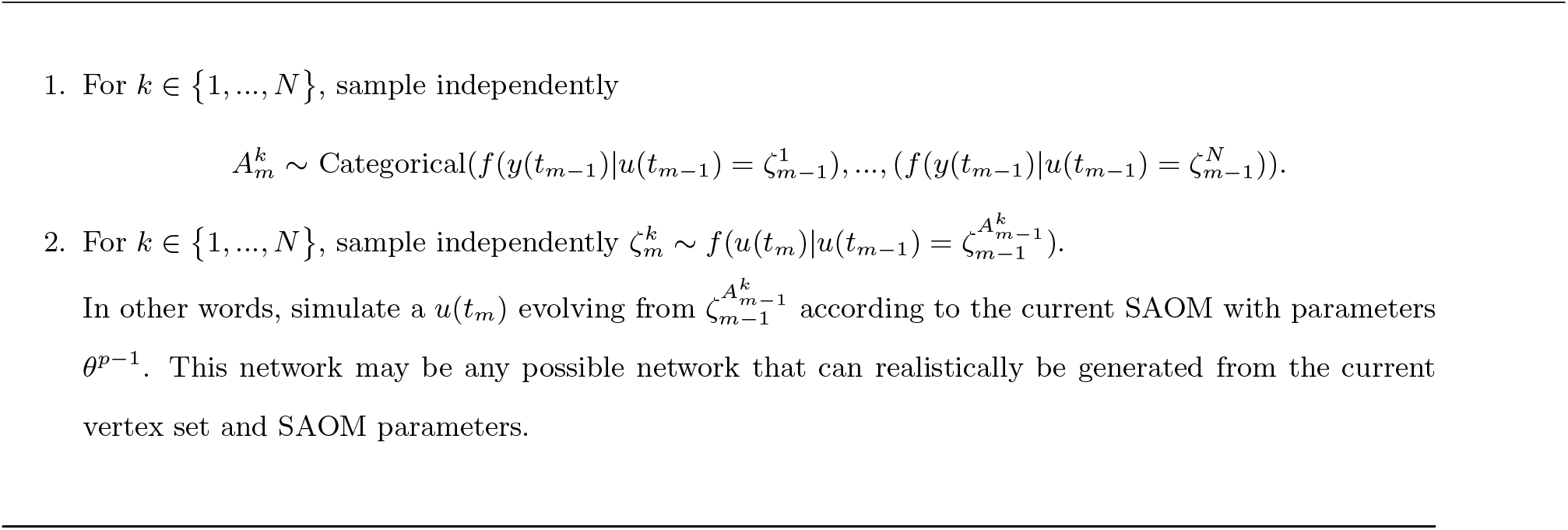

One can view Algorithms 1.1 and 1.2 as a kind of evolutionary system where at observation moment *m < M* each particle has exactly one parent, and at each observation moment *m >* 1, each particle has some number of offspring. Algorithms 1.1 and 1.2 create a collection of possible true networks at each observation moment. Figure 2 provides a visual description of the scheme. However, we ultimately would like a sample of possible true network *series*. Algorithm 1.3 describes how to sample such a series. By sampling an ancestral line, we are effectively sampling from 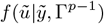.

**Figure 2:**
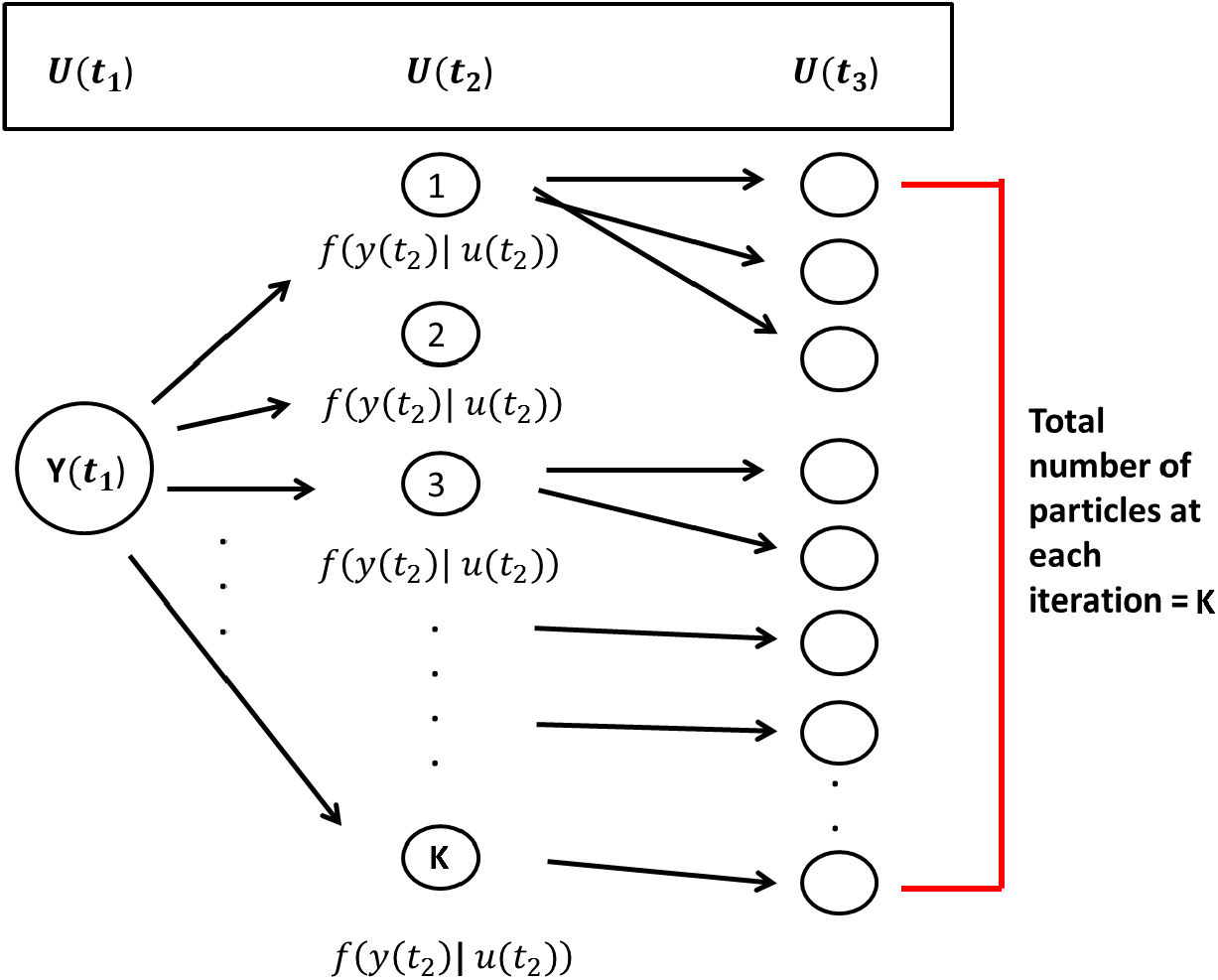
Particle Filtering Sampling Scheme. *K* particles are sampled at each observation moment following a two-stage process. First, particles are selected from the previous observation moment with probability proportional to the conditional distribution of the true network given the observed network at that time point. Then, a true network (i.e., particle) at the next observation moment is sampled/simulated, starting from the current selected particle network, and according to the parameter estimates in the SAOM.

#### Algorithm 1.3

Sampling an ancestral line (i.e., a true network series)

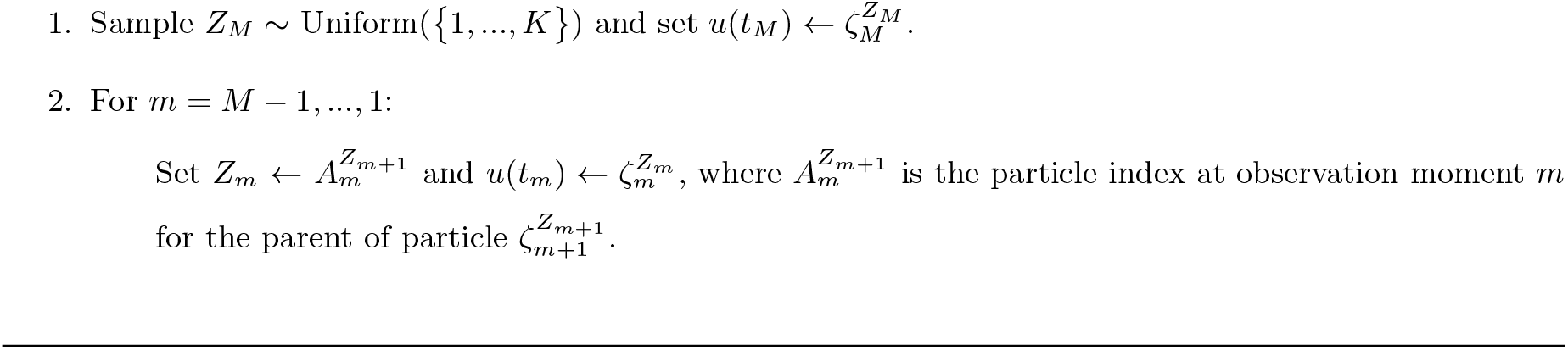

### 4.4 Putting it all together

We now combine the elements presented in the previous sections to define a complete algorithm for the estimation of *α, β*, and *γ* in our SAOM-HMM.

Given a series of observed networks 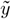 and a number of effects one wishes to include in the SAOM, a SAOM is fit (via maximum likelihood estimation) to get initial estimates of the parameters associated with each effect. We also make the choice, for this current implementation, to assume a constant rate parameter across all vertices. This is a fairly standard choice and is recommended in the SAOM literature, unless one has strong reason to believe otherwise. Set the initial SAOM parameter estimates, 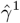, equal to these estimates.

The *p*^*th*^ updating step of our EM algorithm then proceeds as follows:

1. Using 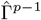, perform Algorithm 1.2 to get *K* possible true networks at each observation moment.
2. Sample *H* number of true network series from an approximation of 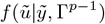 using the ancestral sampling scheme in Algorithm 1.3.
3. Perform the maximization step using Equations 4.3 to obtain 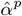 and 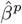. This step obtains estimates of the false positive and false negative rates.
4. Perform the maximization routine outlined in section 4.2.2 to obtain 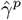. This step provides estimates of the SAOM parameters.
5. Repeat steps 1-4 until convergence. The convergence criteria we have used in our simulation study is based on a moving average and is the following:

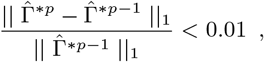

where 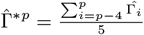and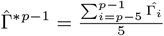.

However, in other instances, some parameter coordinates may have a different scale than others, and the true estimate may be close to 0. Therefore, the convergence criterion may need to be adapted.

The choice of *K* in step 1 (i.e. the number of particles sampled at each observation moment) and the choice of *H* (i.e. the sample of true network series to sample) in step 2 should depend on the size of the network, the number of observation moments, and the amount of noise and strength of the SAOM parameter signals one suspects to be present. For example, when working with network sizes of 10 vertices, 4 observation moments, and 5 parameters in our SAOM, we have used a K of 50,000 and an *H* of 3000 for the maximization of *α* and *β*. For the maximization of *γ*, we have worked with an *H* of 50.

We deliberately keep *H* relatively small (in this case, 50) because this step involves finding 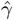 that maximizes data from *H* number of network series. As has often been remarked by the developers of SAOMs, even estimating *γ* for one series of networks can be time consuming, depending on the size of the network and the number of parameters in the model. This step of our algorithm consumes much of the run time, and to try and estimate *γ* for many more *H* will require significantly more time.

### 4.5 Calculation of Standard Errors

The algorithm outlined thus far only produces the parameter estimates. Additional work is required if one wants the standard errors associated with the estimates. Since inference is likely the end goal in practice, a method for calculating standard errors of the estimates is needed. We propose performing a parametric bootstrap where the maximum likelihood estimates from the above algorithm are collected and a number of network series from the estimated model are sampled. The proposed algorithm is then run on each of the sampled network series to obtain a new collection of parameter estimates, which are then used to calculate the standard errors. The more samples we take, the more accurate the estimates of the standard error will be. However, again, there is a computational trade-off, since our algorithm needs to be run for each of the sampled network series.

### 4.6 Algorithm Modifications

The focus of the framework discussed thus far has been on directed networks since SAOMs were initially developed for directed networks. It should be noted, though, that SAOMs are now capable of handling undirected networks, and so is our methodology. The only caveat is that one needs to define how edges are assumed to form in the SAOM model [36]. For example, does one vertex unilaterally impose that an edge is created or dissolved? Or, do both vertices have to ‘agree’? We present an example of an application to undirected networks in Section 6.

Another modification one may choose to make is with regards to how *γ* is estimated in the maximization step in the algorithm we propose. In section 4.2.2, we outline a Robbins Monro algorithm to maximize the complete data log likelihood. Our approach borrows from the algorithm Snijders et al. use for the maximum likelihood estimation of the SAOM parameters for the evolution of one observed network series [35]. The computation for this step is time consuming. As a way to reduce the maximization time for *γ*, one could instead perform a method of moments based estimation routine. This approach also utilizes a Robbins Monro algorithm. In the maximum likelihood based routine, at each iteration of the Robbins Monro algorithm, possible paths leading from one network to the next are augmented for each of our sampled true network series. A complete data score function is then calculated and used in the updating step. The method of moments based routine instead simulates the SAOM evolution process, calculates statistics corresponding to each parameter in the model on both the observed and simulated networks, and then takes the difference of these to be used in the updating step of the Robbins Monro algorithm. In other words, parameter estimates are determined as the parameter value for which the expected value of the statistics equals the observed value at each observation point and for each sampled true network series [37].

Although the method of moment based estimation for *γ* does not produce formal maximum likelihood estimates, it it still a viable estimation method and has been shown to provide similar results in our setting. We demonstrate this in section 5.2 through a small simulation study. In larger networks, the savings in run time may *j*ustify the use of method of moments as an approximation.

### 4.7 Algorithm Implementation

The algorithm outlined in this paper is implemented in R. We have written code for steps 1-3 outlined in Section 4.4. For step 4, we call upon the RSiena package [25], which was developed by R. Ripley, K. Boitmanis, and T.A. Snijders for the implementation of SAOMs. Version 1.1-232 was used for our modeling framework. Minor modifications were made to the source files of this package (saved locally) to allow for our specific algorithm outlined in Section 4.2.2 to be implemented.

One iteration of our E-M algorithm for the networks used in our simulation study explained in the following section takes approximately 45 minutes to run on a high performance Linux computing cluster using 5 cores. The algorithm converged, on average, after 15-20 iterations.

## 5 Simulation Study

### 5.1 Study Design

We present a small simulation study to demonstrate the accuracy of our method and draw a comparison between the behavior of our HMM-SAOM ML estimator and the ML estimator obtained from only fitting a SAOM (i.e. the naive approach). For this study, we simulate 10 node directed networks at 4 observation moments, referred to as *t*_*1*_, *t*_*2*_, *t*_*3*_, and *t*_*4*_. We also create 2 vertex covariates, called Covariate A and Covariate B. Both are indicator variables and are defined as:

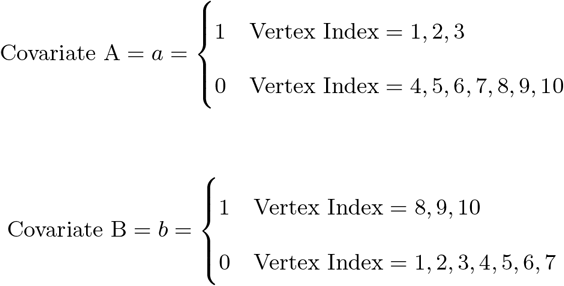

The objective function in the SAOM for the evolution of the true networks, *u*(*t*_*m*_), contains 5 effects (2 endogenous and 3 exogenous). Table 2 lists these effects, their mathematical definitions, and their descriptions. The large, negative value for outdegree, keeps our simulated true networks fairly sparse. It sets the probability of connections forming low, unless the other parameters in the function influence specific vertices in a more positive way. For example, the large positive value for reciprocated edges encourages a directed edge to form if one already exists in the reverse direction. These parameters, in con*j*unction with the parameters assigned to the three covariate related effects, promote the network structure demonstrated in Figure 3. In other words, the Covariate A Ego effect makes it highly likely that vertices 1, 2, and 3 will initiate out-connections to other vertices, outweighing the negative density parameter, as long as these connections form with vertices 8, 9, and 115 since the Covariate B Alter effect parameter is high. Furthermore, the Covariate B Ego effect parameter is large, encouraging vertices 8, 9, and 10 to connect to other vertices, but they will mainly choose to connect amongst each other and to vertices 1, 2, and 3 due to the large reciprocity parameter.

**Table 2:**
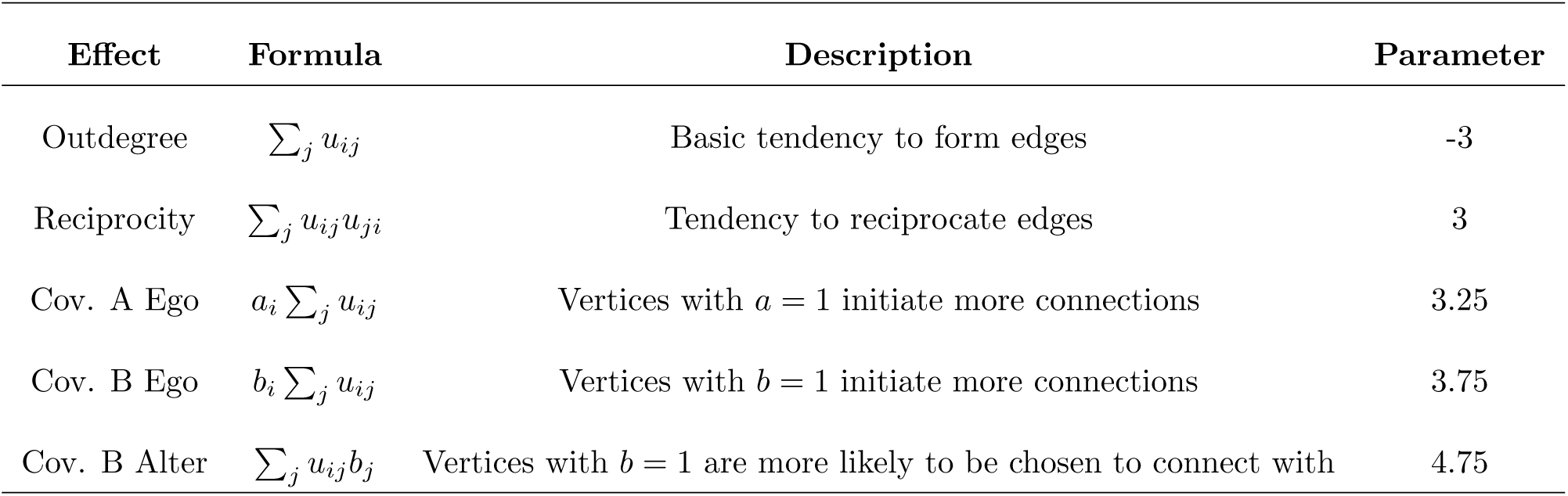
Simulation Model Effects and Parameters.

**Figure 3:**
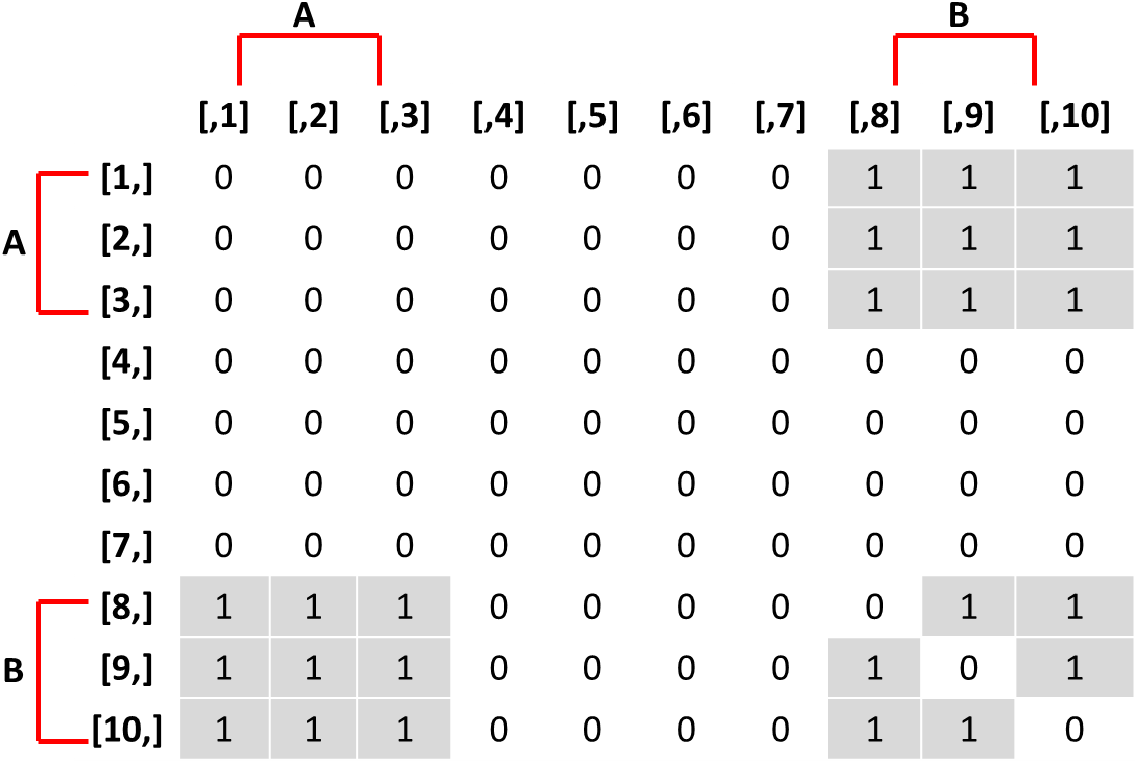
Adjacency matrix of true networks encouraged by the SAOM in the simulation study. An entry of 1 in row *i* and column *j* represents a directed edge from vertex *i* to vertex *j*.

To create an initial true directed network without error at time *t*_*1*_ for the evolution process to begin at, we first created a random network of approximately one quarter of the possible edges. We chose this density to be in line with the density of the encouraged network structure of the simulation model. We then simulated a next network state, evolving from this network, with a small rate parameter of 1 and the true parameter values we set for our SAOM objective function. By doing this, we created a network that had begun to drift towards a network state that has a high probability under the stationary distribution. This network was used as the first observed and true network for every simulation in our study.

We performed 100 simulations each, for 3 different error rate scenarios, while holding the first network constant and keeping the same SAOM objective function parameters. For each simulation, we simulated true networks at the 2^*nd*^, 3^*rd*^, and 4^*th*^ observation moments, according to our true SAOM and with a constant rate parameter of 3. This rate was small enough that the networks gradually approached dynamic equilibrium, thus simulating a realistic evolution process. However, it is large enough that the network at *t*_*4*_ was a network in (or nearly in) dynamic equilibrium. We then created ‘observed’ networks by introducing error to the edges of each true network. True edges remained edges in the observed network with probability equal to 1 − *β* and non-edges in the true networks remained non-edges in the observed network with probability 1 − *α*. We then fit the HMM-SAOM to the observed networks, as well as just the SAOM by itself.

To determine the error-rates, we first ran 200 simulations of our true network evolution process to determine the expected number of edges and non-edges. The expected edges were calculated to be 21.31 and the expected non-edges to be 68.69. Therefore, we chose three error-rate scenarios. We performed 100 simulations with an expected number of 3 false edges and 3 false non-edges, which equates to an *α* of 0.044 and a *β* of 0.141. We performed an additional 100 simulations where we reduced *α* and *β* to half of what we originally defined them to be, giving us *α* = .022 and *β* = .0705, and lastly, we performed 100 simulations where we doubled the original *α* and *β*, equating to *α* = .088 and *β* = .282.

### 5.2 Simulation Study Results

Table 3 reports the average estimates and the standard deviations for each estimate, under each scenario, based on 100 simulations. The root mean squared errors are presented in Table 5, as well as the estimated relative mean squared error (MSE) of the HMM-SAOM estimator compared to the SAOM only estimator (ie. ignoring measurement error). Side-by-side boxplots of the distribution of each parameter estimate are also included. Our results demonstrate that the HMM-SAOM estimator out performs the SAOM-Only estimator under various error rates. The estimated relative MSE for these five SAOM parameters ranges from 0.053 to 0.468, indicating that the HMM-SAOM parameters are much more accurate than the SAOM parameters when we have observed networks with error. The bias observed in our estimators is not unexpected. The same is shown in the maximum likelihood estimates in the standard SAOM framework. When we simulate 100 error-free networks with our true parameters and then estimate the parameters via the standard MLE SAOM routine, we obtain mean estimates that are similar to those obtained under our HMM-SAOM estimator, despite our method needing to account for noise, as well. Table 4 displays these results. Note, however, that the bias shown in our estimator is substantially smaller than that for the SAOM fit to these same noisy data. When we increase our network size to 30 vertices, we still see a great improvement in the SAOM-HMM compared to the standard SAOM (see Supplementary Appendix C for additional details).

**Table 3:**
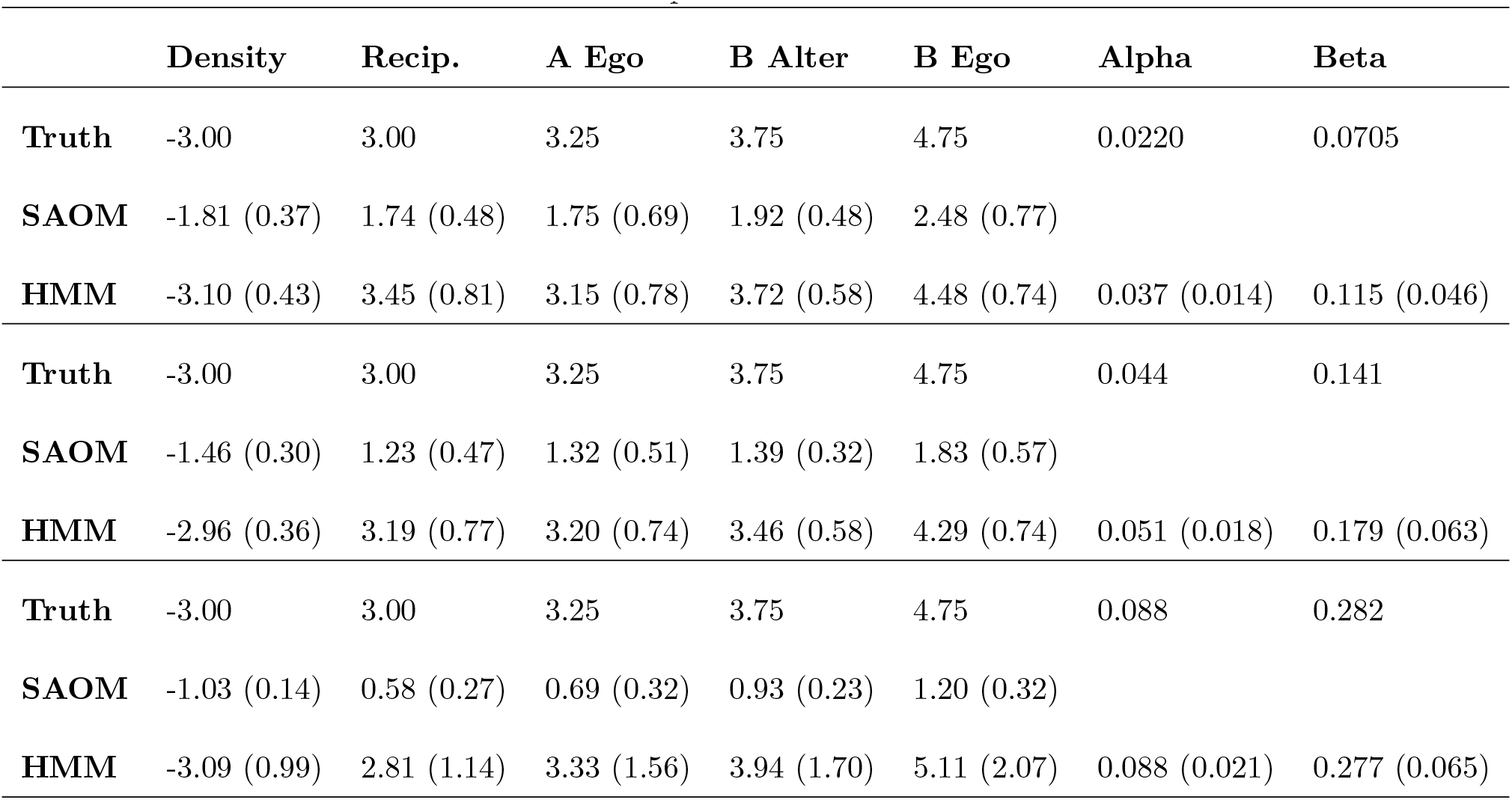
Mean and standard deviation of parameter estimates based on 100 simulations.

|  | Density | Recip. | A Ego | B Alter | B Ego | Alpha | Beta |
| --- | --- | --- | --- | --- | --- | --- | --- |
| **Truth** | -3.00 | 3.00 | 3.25 | 3.75 | 4.75 | 0.0220 | 0.0705 |
| **SAOM** | -1.81 (0.37) | 1.74 (0.48) | 1.75 (0.69) | 1.92 (0.48) | 2.48 (0.77) |  |  |
| **HMM** | -3.10 (0.43) | 3.45 (0.81) | 3.15 (0.78) | 3.72 (0.58) | 4.48 (0.74) | 0.037 (0.014) | 0.115 (0.046) |
| **Truth** | -3.00 | 3.00 | 3.25 | 3.75 | 4.75 | 0.044 | 0.141 |
| **SAOM** | -1.46 (0.30) | 1.23 (0.47) | 1.32 (0.51) | 1.39 (0.32) | 1.83 (0.57) |  |  |
| **HMM** | -2.96 (0.36) | 3.19 (0.77) | 3.20 (0.74) | 3.46 (0.58) | 4.29 (0.74) | 0.051 (0.018) | 0.179 (0.063) |
| **Truth** | -3.00 | 3.00 | 3.25 | 3.75 | 4.75 | 0.088 | 0.282 |
| **SAOM** | -1.03 (0.14) | 0.58 (0.27) | 0.69 (0.32) | 0.93 (0.23) | 1.20 (0.32) |  |  |
| **HMM** | -3.09 (0.99) | 2.81 (1.14) | 3.33 (1.56) | 3.94 (1.70) | 5.11 (2.07) | 0.088 (0.021) | 0.277 (0.065) |

**Table 4:** Mean and standard deviation of parameter estimates based on the standard SAOM MLE routine for 10 node noise-free networks.

|  | Density | Recip. | A Ego | B Alter | B Ego |
| --- | --- | --- | --- | --- | --- |
| **Truth** | -3.00 | 3.00 | 3.25 | 3.75 | 4.75 |
| **Standard SAOM** | -2.85 (0.38) | 2.82 (0.61) | 3.02 (0.74) | 3.50 (0.56) | 4.39 (0.81) |

We see an increase in variability in the estimates for our larger error rate scenario, which is expected. Interestingly, the opposite phenomena appears to be true when only fitting a SAOM. We suspect this is due to the fact that true signal becomes more diluted as more error is introduced, encouraging the parameter estimates to consistently stay right above the 0 mark (i.e there is a consistent weakening of the signal since there a large amount of noise). Whereas, when smaller amounts of error are introduced, which edges end up being affected by the error play a ma*j*or role in the estimates the SAOM model produce. It may be the case that the small amount of noise impacts edges in a way that do not dilute the signal as much.

**Table 5:** Root mean squared error and relative MSE for the SAOM objective function parameter estimates. The SAOM only estimates are the reference group for the relative MSE.

| | $\alpha = 0.022, \beta = 0.0705$ | | | $\alpha = 0.044, \beta = 0.141$ | | | $\alpha = 0.088, \beta = 0.282$ | | |
| --- | --- | --- | --- | --- | --- | --- | --- | --- | --- |
|  | HMM | SAOM | Rel. MSE | HMM | SAOM | Rel. MSE | HMM | SAOM | Rel. MSE |
| **Density** | 0.435 | 1.242 | 0.123 | 0.362 | 1.570 | 0.053 | 0.996 | 1.977 | 0.254 |
| **Reciprocity** | 0.923 | 1.349 | 0.468 | 0.797 | 1.830 | 0.190 | 1.160 | 2.432 | 0.227 |
| **Cov. A Ego** | 0.782 | 1.648 | 0.225 | 0.740 | 2.001 | 0.137 | 1.561 | 2.576 | 0.367 |
| **Cov. B Alter** | 0.581 | 1.891 | 0.094 | 0.650 | 2.379 | 0.075 | 1.708 | 2.829 | 0.365 |
| **Cov. B Ego** | 0.783 | 2.391 | 0.107 | 0.872 | 2.971 | 0.086 | 2.105 | 3.565 | 0.349 |
| **Alpha** | 0.020 |  |  | 0.019 |  |  | 0.021 |  |  |
| **Beta** | 0.064 |  |  | 0.073 |  |  | 0.066 |  |  |

We also perform 100 simulations under our middle error rate scenario where we use the SAOM method of moments estimation routine for *γ* in place of the maximum likelihood estimation routine in our M-step of the E-M algorithm (as discussed in section 4.6). Results are presented in Table 6. We see that substitution of the method-of-moments approach within our algorithm yields results similar to the originally described algorithm in the sense that the absolute bias is not much larger. However, several of the parameters have bias in the opposite direction. There is also some increase in variability of the estimates. We suspect that the method-of-moments approach will produce estimates that are more accurate than the ones displayed in Table 6 for larger network sizes. Despite the increase in variability observed when performing the method-of-moments estimation routine for *γ*, it may be worthwhile to use when working with larger networks or many observation moments.

**Table 6:** Mean and standard deviation of parameter estimates for MLE vs. MoM for estimation of *γ*.

|  |  | MLE |  | MoM |  |
| --- | --- | --- | --- | --- | --- |
|  |  | # of Sims = 100 |  | # of Sims = 100 |  |
|  | Truth | SAOM | HMM | SAOM | HMM |
| **Density** | -3.00 | -1.46 (0.30) | -2.96 (0.36) | -1.43 (0.31) | -3.33 (0.64) |
| **Reciprocity** | 3.00 | 1.23 (0.47) | 3.19 (0.77) | 1.21 (0.40) | 3.52 (0.97) |
| **Cov. A Ego** | 3.25 | 1.32 (0.51) | 3.20 (0.74) | 1.38 (0.59) | 3.54 (1.22) |
| **Cov. B Alter** | 3.75 | 1.39 (0.32) | 3.46 (0.58) | 1.39 (0.36) | 4.07 (1.05) |
| **Cov. B Ego** | 4.75 | 1.83 (0.57) | 4.29 (0.74) | 1.95 (0.60) | 5.35 (1.19) |
| **Alpha** | 0.044 |  | 0.051 (0.018) |  | 0.055 (0.019) |
| **Beta** | 0.141 |  | 0.179 (0.063) |  | 0.166 (0.062) |

**Figure 4:**
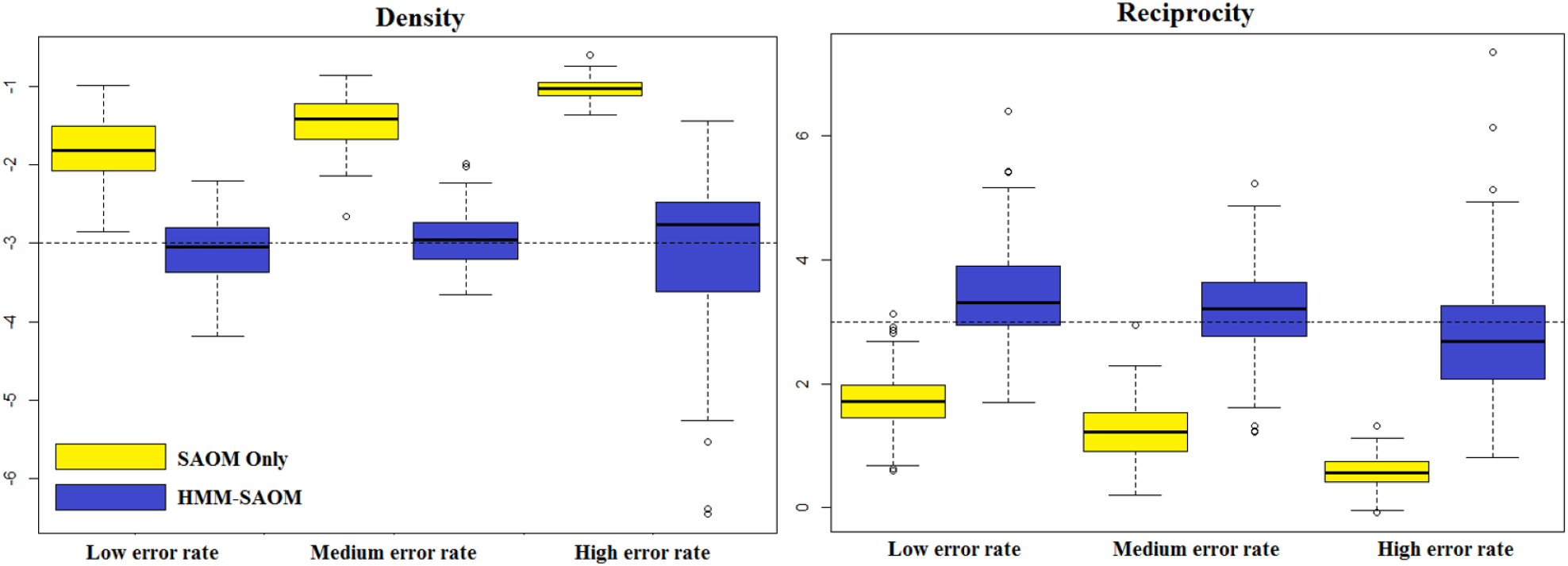
Boxplots for Density and Reciprocity Parameter Estimate Distributions obtained from 100 simulations for our HMM-SAOM model and also for the SAOM only (i.e. the naive approach). The dashed line represents the true parameter value.

**Figure 5:**
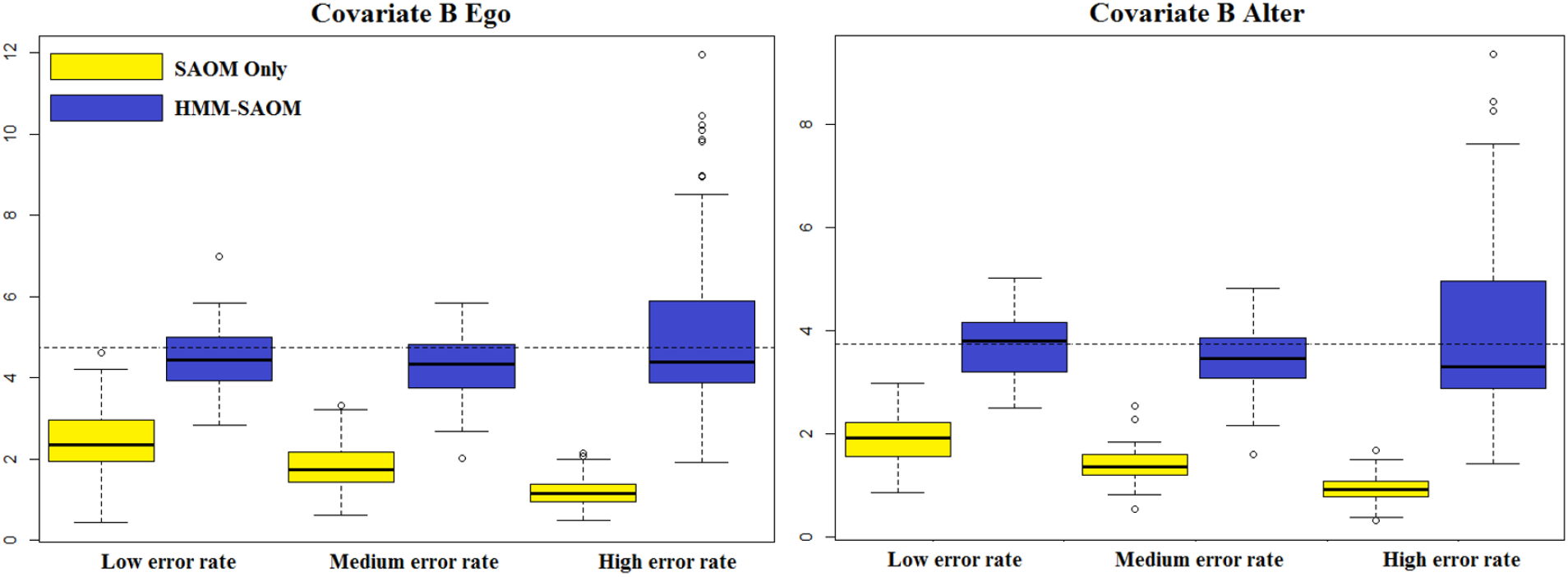
Boxplots for Covariate B Alter and Ego Parameter Estimate Distributions obtained from 100 simulations for our HMM-SAOM model and also for the SAOM only (i.e. the naive approach). The dashed line represents the true parameter value.

**Figure 6:**
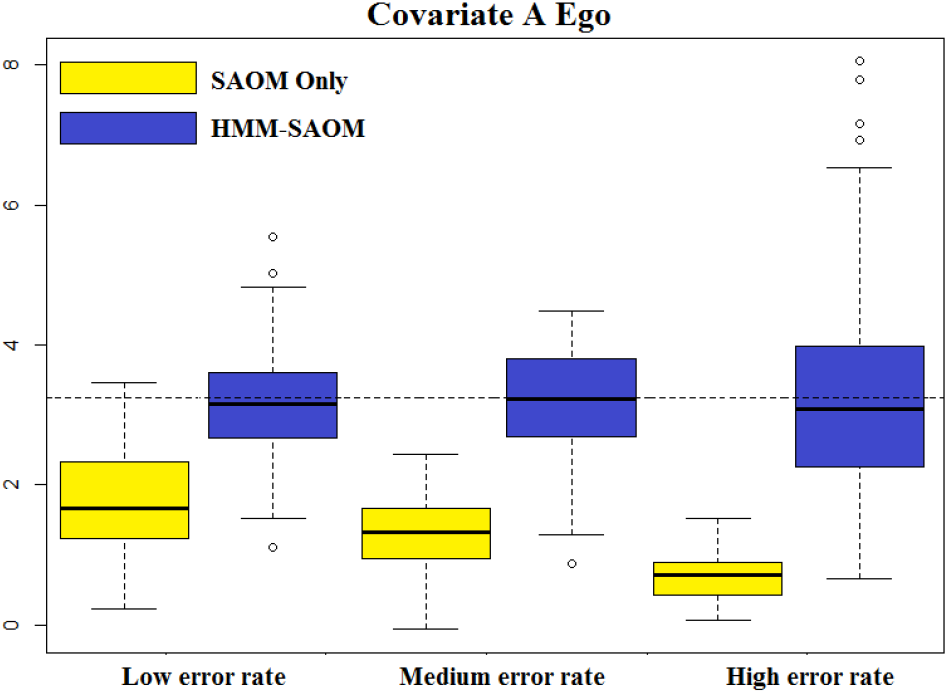
Boxplots for Covariate A Ego Parameter Estimate Distributions obtained from 100 simulations for our HMM-SAOM model and also for the SAOM only (i.e. the naive approach). The dashed line represents the true parameter value.

## 6 Analysis of EEG Complex Functional Networks

We apply this methodology in an analysis of dynamic EEG complex functional networks. Networks are becoming a popular model to illustrate both the physiological connections (structural networks) and the coupling of dynamic brain activity (functional networks) linking different areas of the brain [1]. Deviations in the expected behavior and organization of brain networks have been observed in several diseases [19, 42, 40, 30]. Although physiological coupling between brain regions must be mediated by underlying structural connectivity, previous studies have found little relationship between large scale structural and functional brain networks [3, 22, 21, 18]. We hypothesize that this may be due to the inherent false positive and false negative error rates present in inferring functional brain networks using current techniques, and therefore that the effect size of structural connectivity and other observed parameters will be increased using our approach. We explored this relationship by relating functional networks inferred from statistical associations between source imaging of EEG activity and underlying cortico-cortical structural brain connectivity determined by probabilistic white matter tractography.

A patient with high density EEG (70 electrodes), digitized electrode coordinates, and high resolution diffusion tensor imaging (DTI; 60 diffusion-encoding directions, 1.85 mm isotropic voxels) was retrospectively identified from clinical evaluations performed at the Massachusetts General Hospital Athinoula A. Martinos Center for Biomedical Imaging between 1/2009 and 12/2012, where she was undergoing evaluation due to epilepsy. The EEG was recorded with a 70-channel electrode cap, based on the 10-10 electrode-placement system (Easycap, Vectorview, ElektaNeuromag, Helsinki, Finland) in the quiet resting state. The positions of the EEG sensors were determined prior to data acquisition with a 3D digitizer (Fastrak, Polhemus Inc., Colchester, VA). The sampling rate was 600 Hz and the data were filtered with high- and low-pass filters from 1-50 Hz for analysis using the MATLAB Signal Processing Toolbox and custom software. Source analysis of EEG data was performed using the MNE software package [14] with anatomical surfaces reconstructed using Freesurfer [10] with 70 vertices distributed across the cortical surface.

Functional undirected binary networks based on the source EEG data were constructed for each contiguous 1 second interval, using cross-correlation as the measure of coupling, as described in Chu et al. [3]. For our analysis, binary networks were averaged across 10 s segments to create a representative weighted network reflecting the average properties of the functional networks over time, where the edge weight or strength reveals the consistency of an edge appearance across time. We then binarized these weighted matrices to obtain a single 70 node, binary network for 4 consecutive 10 s intervals. The networks were binarized by choosing a threshold for each network that kept the density at approximately 16% − 18%. To infer structural connectivity networks, probabilistic tractography (Probtrackx2 through FSL 5.0.4/FDT-FMRIB’s Diffusion Toolbox 3.0; FMRIB’s Software Library) was used to process the DTI data obtained from a 3 T Magnetom Trio scanner, choosing as seed and target regions of interest (ROIs) the same cortical vertices used to infer functional networks, and a weighted structural connectivity matrix was constructed. Please see Chu et al. for a more detailed description of data acquisition and network construction [3].

Since we are working with undirected networks, we will work under the assumption that one ROI takes the initiative and unilaterally imposes that an edge is created or dissolved. Table 7 lists the effects we chose to place into the objective function of our SAOM. We would like to test hypotheses involving how each of these effects drives change in this individual’s functional EEG networks. Moreover, we would like to investigate which of these effects are detectable in HMM-SAOM approach, but not the SAOM alone. A short description of these effects is given below.

### Endogenous Effects

#### Density

Represents the basic tendency for vertices to form connections. It is similar to an intercept in a regression model. For sparser networks, this parameter will often be negative.

#### Transitive Triads

Represents the tendency for vertices to form connections that position them within triangular structures. Triangles serve as a representation of clustering and clustering serves as an indicator of segregation within the network. When examining functional connectivity data in healthy individuals, network analysis has revealed a notable high clustering coefficient. This coefficient is linked to elevated local efficiency in information transfer, specifically facilitating specialized processing [28]. Triangles also hold significance from a motif perspective. Network motifs are intriguing in functional brain networks as they signify distinct topological connection patterns, essentially serving as the“building blocks” of the entire network [39, 38].

#### Number of Vertices at Distance Two

Defined by the number of ROIs to whom *i* is indirectly connected (through at least one intermediary). When this effect has a negative parameter, vertices will have a preference for having few others at a geodesic distance of 2. Short path lengths play a crucial role in fostering functional integration and efficiency by facilitating communication with minimal intermediate steps, thereby reducing the impact of noise or signal degradation [38]. Research has demonstrated that functional networks in diverse brain disorders exhibit longer path lengths, suggesting a less efficient organization of connectivity [27, 45].

### Exogenous Effects

#### Electrode Distance

Represents the tendency for vertex pairs with higher values of electrode distance to form connections. A negative parameter implies that vertex pairs with a larger distance separating them, have a smaller probability of connecting.

#### Structural Connectivity

Reflects the tendency for vertex pairs with higher values of structural connectivity to form connections. A positive parameter implies that vertex pairs with more structural connectivity, have a higher probability of connecting.

**Table 7:**
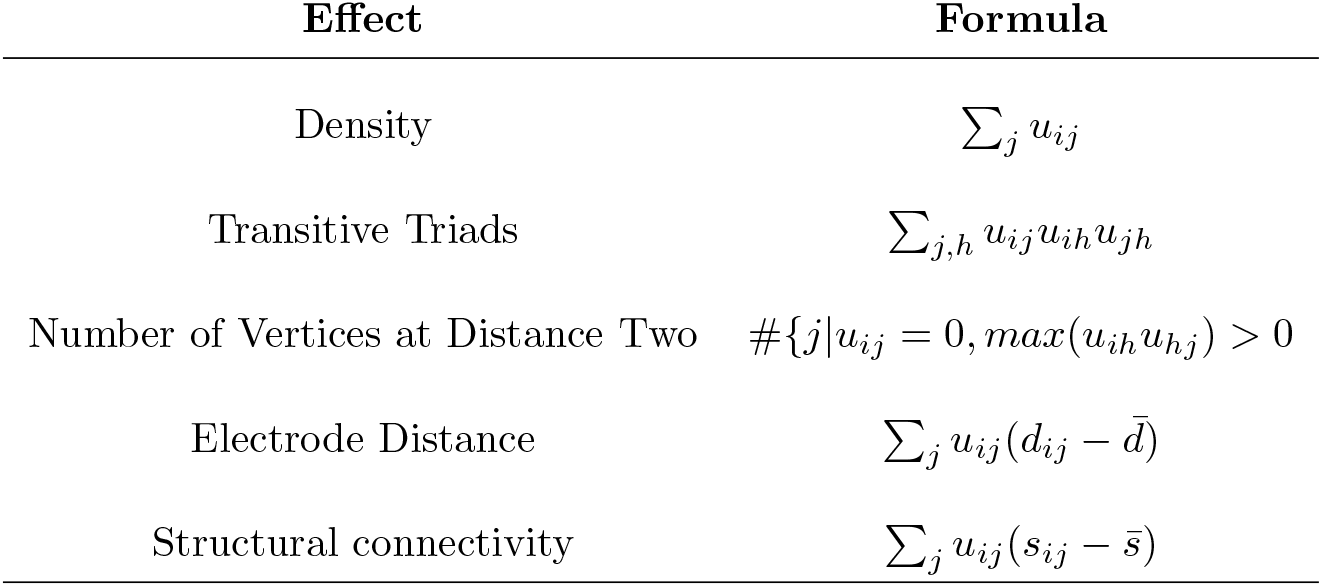
Mathematical Definition of SAOM Effects for EEG Functional Network Analysis.

| Effect | Formula |
| --- | --- |
| Density | $\sum_j u_{ij}$ |
| Transitive Triads | $\sum_{j,h} u_{ij} u_{ih} u_{jh}$ |
| Number of Vertices at Distance Two | $\#\{j \vert u_{ij} = 0, \max(u_{ih} u_{hj}) > 0\}$ |
| Electrode Distance | $\sum_j u_{ij} (d_{ij} - \bar{d})$ |
| Structural connectivity | $\sum_j u_{ij} (s_{ij} - \bar{s})$ |

We first fit a SAOM to our series of 4 observed functional networks. We then fit our HMM-SAOM to the same series of functional networks. Standard errors for the HMM-SAOM were approximated via parametric bootstrap in the manner described in section 4.5 with 10 sampled network series. The estimated parameters, standard errors, and t-ratios for both models are reported in Table 8. We must interpret the results cautiously, based on the relative magnitude of t-ratios, as formal grounds for comparison to the t distribution are not well established. It is also important to note that the parameter estimates allow for a caricature of the rules governing the dynamic change in the network [41]. Because the temporal progression is taken care of by the rate functions, the parameters in the objective function are static and are comparable across periods of different lengths of time. As the SAOM authors point out [41], a common misunderstanding is that the parameter estimates express tendencies over time. Instead, they should be interpreted as satisfaction measures that are suitable for explaining the observed changes.

**Table 8:** EEG Functional Network Analysis Results from fitting our HMM-SAOM and from only fitting a SAOM.

|  | SAOM Only |  |  | HMM-SAOM |  |  |
| --- | --- | --- | --- | --- | --- | --- |
|  | Estimate | S.E. | T-Ratio | Estimate | S.E. | T-Ratio |
| **Density** | -1.437 | 0.044 | -32.659 | -2.485 | 0.154 | -16.136 |
| **Transitive Triads** | 0.094 | 0.003 | 31.333 | 0.161 | 0.014 | 11.500 |
| **Number at Distances 2** | -0.082 | 0.016 | -5.125 | -0.034 | 0.167 | -0.204 |
| **Electrode Distance** | -0.014 | 0.001 | -14.000 | -0.022 | 0.002 | -11.00 |
| **Structural Connectivity** | 3.596 | 0.668 | 5.383 | 6.991 | 1.777 | 3.934 |
| **Alpha** |  |  |  | 0.135 | 0.008 | 16.875 |
| **Beta** |  |  |  | 0.190 | 0.017 | 11.176 |

As hypothesized, we find a positive estimate (and a large t-ratio) for the structural connectivity effect in both models. This indicates that regions with increased structural connectivity are more likely to form functional connections. The difference between the estimates produced by the two models for this effect is quite large. We conclude that the true signal for this effect is strong and may be diluted by measurement error on the network edges. The false positive rate is estimated to be 13.5%, while the false negative rate is estimated to be 19%. Furthermore, as one would expect, the electrode distance effect is associated with a negative parameter estimate, suggesting that the further apart two regions are, the less likely they are to connect. The estimate produced by the SAOM-HMM is approximately 1.57 times the magnitude of that estimated by the SAOM. In addition, the parameter estimate associated with the transitive triads effect is positive and 1.7 times larger in the HMM-SAOM model. The t-ratio is also very large. This indicates that triangle formation is favored as the network evolves. The distance 2 parameter estimate is negative in both models. However, it is half the magnitude and has a much smaller t-ratio in the HMM-SAOM, suggesting a weaker signal than what would have been inferred in the naive method of fitting *j*ust a SAOM.

## 7 Discussion

Our framework shows an increase of MSE over the naive approach of only fitting a SAOM when noise is present in network edges. Therefore, we feel it is a more appropriate modeling choice if error is believed to be present. In our fitting of the HMM-SAOM on EEG functional networks, we find stronger effect sizes in the transitive triads, electrode distance, and structural connectivity effects than what is found when fitting the standard SAOM to the networks. We also obtain an estimate of the false positive and false negeative rates on edge status of the inferred networks, which we would not have obtained by simply fitting the standard SAOM.

As mentioned in the Introduction Section, we take a similar approach to Lospinoso [20], but instead of using a full MCMC algorithm for performing maximum likelihood estimation of the model parameters, we took an MCMC within an E-M approach to parameter estimation. Both approaches have their limitations. In either case, there may be multimodality of the likelihood, potentially causing the E-M to converge to local maxima or for there to be poor mixing of MCMC samplers. Additionally, Lospinoso proposes three potential models for the measurement model, while we focus on one measurement model. Our framework is certainly able to handle more complicated measurement models, but we focused on a model that enumerates false positive and false negative edges in a very simple case given the popularity of quantifying false positive and false negative edges in the brain network science community. One may easily extend our model to another measurement model framework as long as the model parameters can be estimated in closed form. Moreover, the SAOM building block portion of our framework may be exchanged for another dynamic network model abiding by the Markov chain properties, such as the Temporal Exponential Random Graph Model (TERGM) [17]. To do so, one would need to incorporate techniques for MCMC sampling in that setting with our overall framework to enable approximate maximum likelihood estimation. We leave this for future work.

Additionally, in order to perform the particle filtering algorithm, we made an assumption of an error-free first network in the sequence of observed networks. To explore how violations of this assumption affect our results, we performed a small simulation study. We repeated the simulation study described in Section 5.1 of the manuscript (for the low error rates), but with error on the network edges for our first observed network, and we did not find a reduction in model performance. What we anticipate is that error on the first observed network will not have a ma*j*or impact on the performance of our algorithm, as long as there are at least 4 networks in the sequence of observed networks. Having less than 4 networks would decrease the sample size, hurting the algorithm’s ability to accurately estimate parameters, especially with a violation of the error-free assumption on the first network.

Just as Lospinoso reported in his dissertation, and as previously mentioned in this section, we found that higher false positive rates shrink the parameter estimates to 0, falsely suggesting no significant effect on network change. Unlike Lospinoso, though, we did not study the effect of having a non-zero false positive rate (or false negative rate) while holding the other error rate constant at 0. This was simply because in the brain network application, we typically expect there to be comparable levels of false positive and false negative edges.

Lastly, in his dissertation, Lospinoso focuses on the application of his methodology to social networks, whereas we focus on a brain network application. While the entire SAOM framework was developed for the social network setting, we feel extending the models to brain network science is justified. Given existing literature that indicates brain regions often change functional connections as a compensatory mechanism [13, 9, 32] or that ‘hub’ regions shift which regions they communicate with based on instructions for the task at hand [4], an ‘actor-oriented’ or ‘node-oriented’ approach is well-motivated and appealing. With that being said, obviously model interpretation with regards to ‘actor’ preferences and decision-making should not be interpreted too literally. The key is that the model expresses tendencies for connections to form based on each brain region’s current connections/network, its own properties, and the properties of all other brain regions.

Stochastic Actor Oriented Models offer great flexibility in that they are able to represent network dynamics as being driven by many factors/influences. Furthermore, the models allow for the accounting of several different explanations of network change, which may be competing and even complementary. This allows for the testing of effects driving the changes, while controlling for other factors, which better enables researchers to delve in, disentangle, and identify which mechanisms are playing a role (as opposed to focusing on different network characteristics individually). We feel that our contribution complements this framework well and is important, especially for network data that is known to contain large amounts of noise. When this is the case, signal becomes so diluted that the naive approach of fitting only a SAOM, gives very inaccurate estimates. Our HMM-SAOM method gives much more accurate estimates of the SAOM parameters, while also providing estimates of the false positive and false negative rates. SAOMs are already very prominent in the social network literature, but we feel this extension to an HMM setting may potentially spark interest in other research fields (e.g. neuroscience) where noisy data is much more of a concern.

## 8 Acknowledgments

This work was supported by NIH awards 1R01NS095369-01 and K25-EB032903-01, Canadian NSERC RGPIN-21523-153566 and NSERC DGDND-2023-03566, and by the National Institute of General Medical (NIGMS) Interdisciplinary Training Grant for Biostatisticians (T32 GM74905). The authors would like to thank Steven M. Stufflebeam, MD for acquisition of the MRI and EEG data used to generate the human functional and structural brain networks. The computational work reported in this paper was performed on the Shared Computing Cluster which is administered by Boston *U*niversity’s Research Computing Services. (www.bu.edu/tech/support/research/).

## Appendix

### A Derivation for 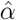 and 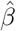

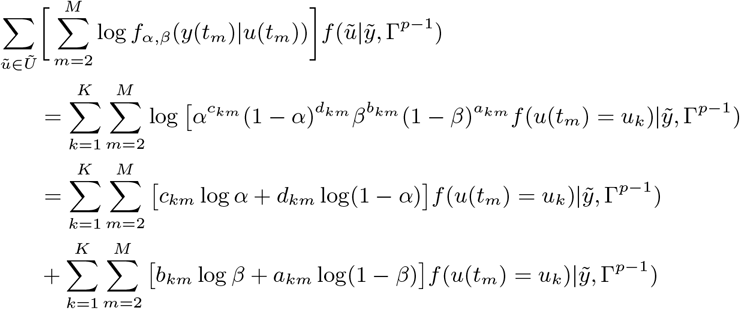

Now, we differentiate with respect to *α*, set the derivative equal to 0, and solve for *α* to get the maximum.

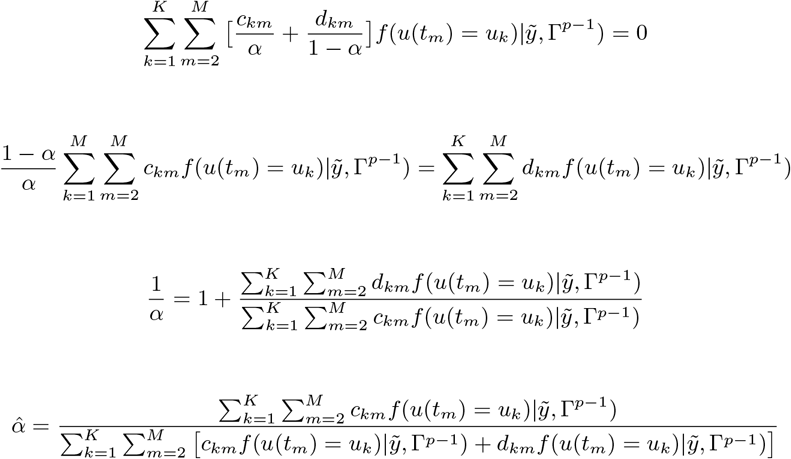

The calculation for 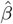 is similar.

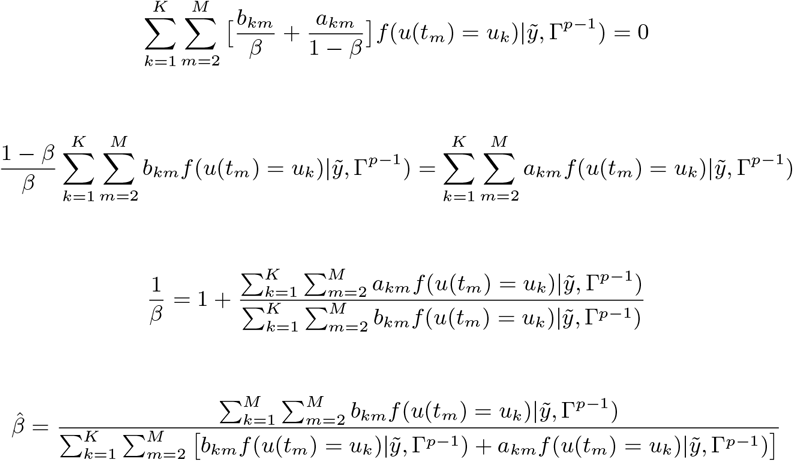

### B Proof of Missing Information Principle in 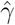 Derivation

Recall that we denote each possible unique true network series by *ũ*. We define the random variable *V* to be a sample path associated with a true network series *ũ*. We prove below that *E*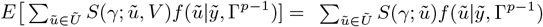. By calculating this expectation, we are able to maximize 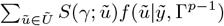 which would otherwise be intractable.

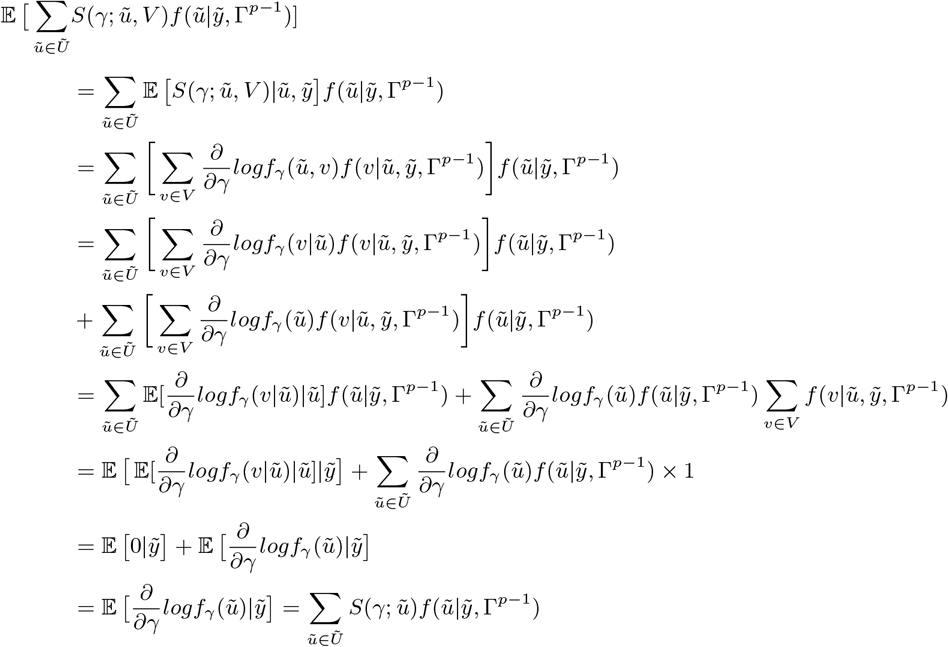

### C 30 Node Network Simulation Study

We present a small simulation study to demonstrate the accuracy of our method on slightly larger network sizes. For this study, we simulate 30 node directed networks at 4 observation moments, referred to as *t*_*1*_, *t*_*2*_, *t*_*3*_, and *t*_*4*_. We also create 2 vertex covariates, called Covariate A and Covariate B. Both are indicator variables and are defined as:

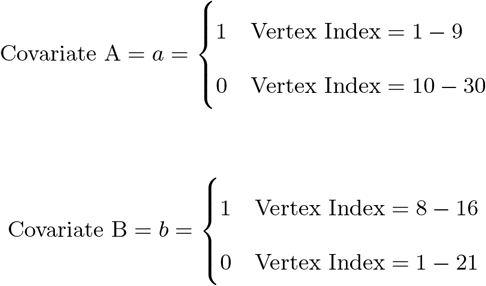

The objective function in the SAOM for the evolution of the true networks, *u*(*t*_*m*_), contains 5 effects (2 endogenous and 3 exogenous). Table 1 lists these effects, their mathematical definitions, and their descriptions. The large, negative value for outdegree, keeps our simulated true networks fairly sparse. It sets the probability of connections forming low, unless the other parameters in the function influence specific vertices in a more positive way. For example, the large positive value for reciprocated edges encourages a directed edge to form if one already exists in the reverse direction. These parameters, in con*j*unction with the parameters assigned to the three covariate related effects, promote the network structure demonstrated in Figure 3 in the manuscript, but with 30 nodes as opposed to 10 nodes and 9 × 9 block sizes, as opposed to 3 × 3 block sizes.

**Table 1:**
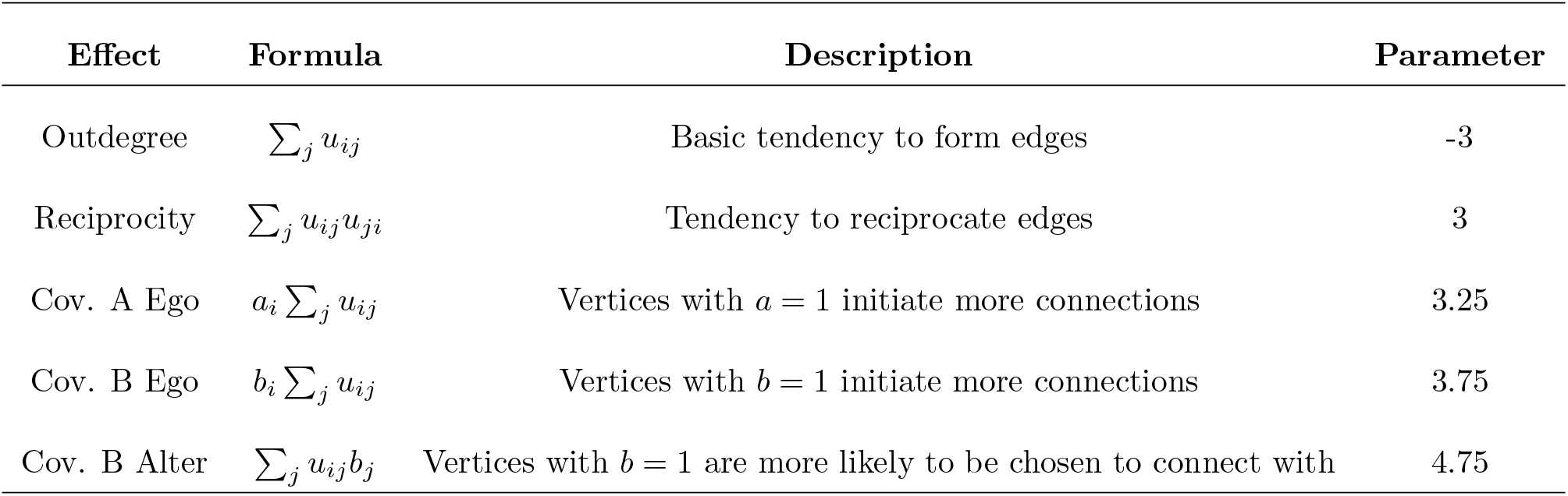
Simulation Model Effects and Parameters.

We performed 315 simulations for the smallest error rate scenario in our paper, while holding the first network constant and the SAOM objective function parameters constant. For each simulation, we simulated true networks at the 2^*nd*^, 3^*rd*^, and 4^*th*^ observation moments, according to our true SAOM and with a constant rate parameter of 3. This rate was small enough that the networks gradually approached dynamic equilibrium, thus simulating a realistic evolution process. However, it is large enough that the network at *t*_*4*_ was a network in (or nearly in) dynamic equilibrium. We then created ‘observed’ networks by introducing error to the edges of each true network. True edges remained edges in the observed network with probability equal to 1-*β* and non-edges in the true networks remained non-edges in the observed network with probability 1-*α*. We then fit the HMM-SAOM to the observed networks.

Table 2 reports the average estimates and the standard deviations for each estimate based on 30 simulations. Our results demonstrate that the HMM-SAOM estimator out performs the SAOMOnly estimator, *j*ust as it did in the 115 node networks presented in the main manuscript.

**Table 2:** Mean and standard deviation of parameter estimates based on 30 simulations.

|  | Density | Recip. | A Ego | B Alter | B Ego | Alpha | Beta |
| --- | --- | --- | --- | --- | --- | --- | --- |
| **Truth** | -3.00 | 3.00 | 3.25 | 3.75 | 4.75 | 0.0220 | 0.0705 |
| **SAOM** | -1.30 (0.08) | 1.28 (0.09) | 1.08 (0.16) | 1.32 (0.09) | 1.61 (0.19) |  |  |
| **HMM** | -3.37 (0.37) | 3.18 (0.48) | 3.55 (0.88) | 3.75 (0.31) | 5.25 (0.77) | 0.058 (0.007) | 0.148 (0.035) |

### D Model Identifiability

#### Theorem 1.

*Let the objective function for the SAOM only contain a parameter for density. In other words, let fi*(*θ*_*1*_, *u*^(*r* − 1)^, *u*^(*r*)^) = *θ*_1_ Σ_*j*_*x*_*ij*_. *Also assume the assumptions of Section 2 hold. Then*, 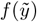 *is globally identifiable*.

*Proof*. We will first show that two sets of parameters where *α*_*2*_ = 1 − *β*_*1*_, *β*_*2*_ = 1 − *α*_*1*_, *θ*_*2*_ = − *θ*_*1*_, and *f*(*U*(*t*_1_) = *u*) = *f*(*U*(*t*_1_) = *ú*) will give the same likelihood for 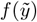 and for *M* = 2, making the model nonidentifiable. We are not assuming assumption 3 in Section 2 is true in this general case. We will then assert that the same holds for *M* of any length and that this is the only parameterization that will cause non-identifiability. Therefore, since our models are assuming *f*(*U*(*t*_1_)) is given (i.e. assumption 3), *f*(*U*(*t*_1_) = *u*) ≠ *f*(*U*(*t*_1_) = *ú*) and the model is identifiable.

We assume that we have two observed network variables, denoted by *Y* (*t*_1_) and *Y* (*t*_2_). The vector with elements *Y* (*t*_1_) and *Y* (*t*_2_), is denoted by 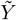. The vector of true network variables, *U*(*t*_1_) and *U*(*t*_2_), underlying the observed networks, is denoted by *Ũ*. Therefore, the likelihood becomes:

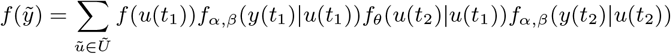

The sum is over all possible true networks series, with a total of 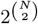 possible networks at each observation time point, where *N* is the number of vertices.

It is trivial to show that for any potential true network *U*(*t*_*m*_) = *u* with *c* edges and *d* non-edges, there is another potential true network *U*(*t*_*m*_) = *ú* with *c* non-edges in place of the *c* edges in *u* and *d* edges in place of the *d* non-edges in *u*. In other words, all of the edges are flipped to non-edges, and all of the non-edges are flipped to edges. Therefore, if given *U*(*t*_*m*_) = *ú, Y* (*t*_*m*_) contains *a* false edges and *b* true non-edges, then given *u*(*t*_*m*_) = *ú, Y* (*t*_*m*_) will contain *a* true edges and *b* false non-edges. Similarly, if given *U*(*t*_*m*_) = *u, Y* (*t*_*m*_) contains *c* false non-edges and *d* true edges, then given *U*(*t*_*m*_) = *ú, Y* (*t*_*m*_) will contain *c* true non-edges and *d* false edges. This means that for each possible *U*(*t*_*m*_) and corresponding 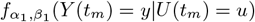, there is a network *ú* where 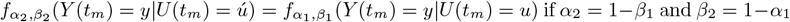.

Next, let’s take each *f*(*U*(*t*_1_)) = *ú* under the second parameterization and set it equal to *f*(*U*(*t*_1_)) = *u* (i.e. the initial distribution of its ‘opposite’ network) under the first parameterization. Now, if we can find a *θ*_*2*_ in which 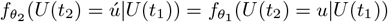then, we have shown that

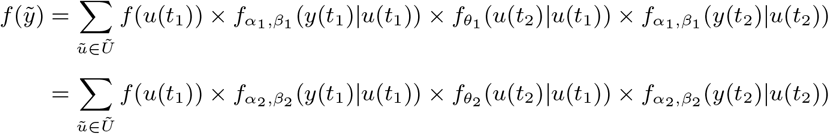

since each summand under the first parameterization will have one, and only one, match to a summand under the second parameterization.

To find *θ*_*2*_ for which 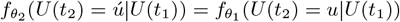, let’s first note that *U*(*t*_*2*_) = *ú* and *U*(*t*_*1*_) = *ú* are the exact opposite networks from *U*(*t*_*2*_) = *u* and *U*(*t*_*1*_) = *u* respectively, in the sense that the edges and non-edges are flipped. Also, recall

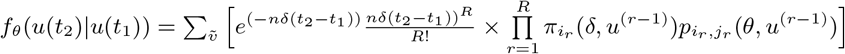

where the sum is over all possible sample paths connecting *u*(*t*_*1*_) and *u*(*t*_2_). If we fix the rate parameter *δ*, then it is sufficient to show that 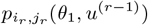 when *U*(*t*_2_) = *u* is equal to 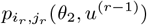 when *U*(*t*_2_) = *ú* for a proof of 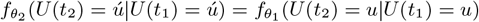.

Let’s first write out the form of *p*_*i, j*_ (*θ*_*1*_, *u*^(*r* − 1)^) for the case when *U*(*t*_2_) = *u*. Recall that:

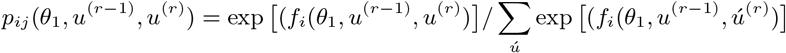

where the summation extends over all possible next network states *ú* and *f*_*i*_(*θ*_*1*_, *u*^(*r* − 1)^, *u*^(*r*)^) only contains an effect for density, and is therefore just equal to *θ*_*1*_ *j* Σ*x*_*ij*_.

Therefore, when it is vertex *i*’s opportunity to make a change in edge status, and if *i* is currently connected to *c* other vertices and has *d* vertices in which it is not connected to (where vertex *j* = *k* is among these), then the probability of *i* forming a connection to *j* = *k* is:

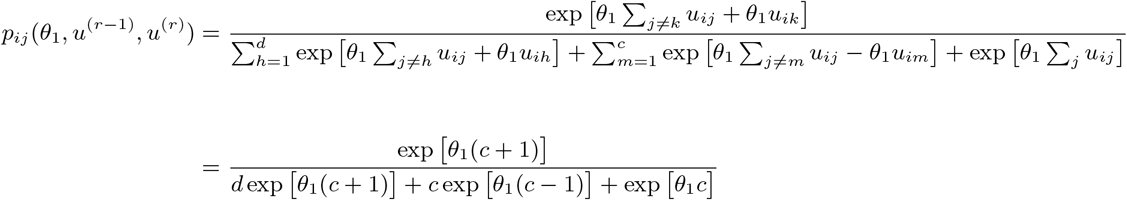

The first term in the denominator is accounting for the fact that vertex *i* could choose to connect to any of the *d* vertices in which it is not connected to. The second term accounts for the fact that vertex *i* could choose to disconnect from any of the *c* vertices in which it is already connected to. And, the last term represents the option of no change.

Now, we’ll move to the case where *U*(*t*_2_) = *ú*. Recall that in this network, all of the edge statuses are opposite to what they were in *U*(*t*_2_) = *u*. Therefore, we want the probability of vertex *i* NOT forming a connection to *j* = *k* to be equal to our prior probability of *i* forming a connection to *j* = *k*. So, when it is vertex *i*’s opportunity to make a change in edge status, and if vertex *i* is currently connected to *d* other vertices (instead of the *c* in it’s ‘opposite’ network), where vertex *j* = *k* is among these, and if vertex *i* has *c* vertices in which it is not connected to (instead of *d*) then the probability of vertex *i* not forming a connection to *j* = *k* is:

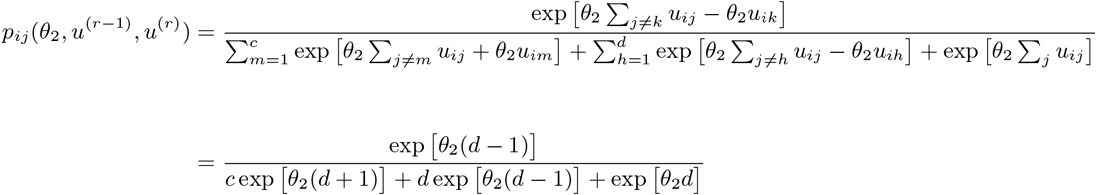

Setting the two probabilities equal to one another gives us…

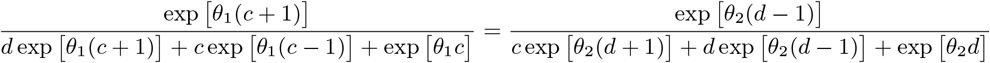

After cross-multiplying we have…

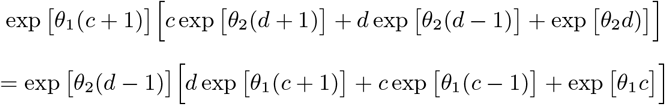

Simplifying the terms to the left of the equal sign gives us:

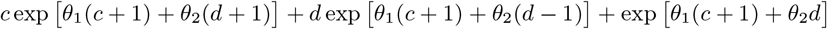

And, simplifying to the right of the equal sign…

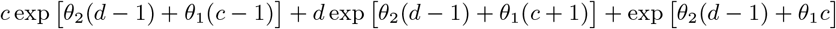

Therefore, for both expressions to equal one another, the following must hold:

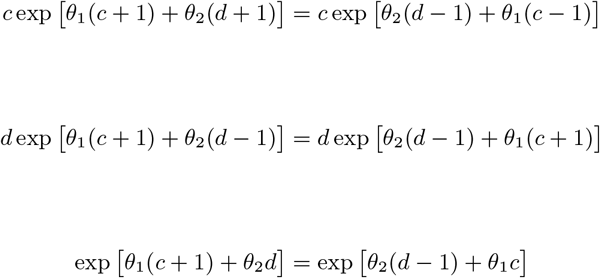

which implies 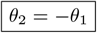.

We have just shown that two sets of parameters (*α*_*1*_, *β*_*1*_, *θ*_*1*_, *f*(*U*(*t*_*m*_) = *u*)) and (*α*_*2*_ = 1 − *β*_*1*_, *β*_*2*_ = 1 − *α*_*1*_, *θ*_*2*_ = − *θ*_*1*_, *f*(*U*(*t*_*m*_) = *ú*) = *f*(*U*(*t*_*m*_) = *u*)) will give the same likelihood for an observed network series where *M* = 2. This is easily extendable to *M* of any length since every possible true network series *Ũ* = *ũ* will have a matching true network series 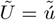 (in which the edge statuses in each network are flipped), where the two have equal probabilities under the two parameterizations explained above.

